# Towards Preclinical Validation of Arbaclofen (R-baclofen) Treatment for 16p11.2 Deletion Syndrome

**DOI:** 10.1101/2023.05.01.538987

**Authors:** Brigitta B. Gundersen, William T. O’Brien, Melanie D. Schaffler, Maria N. Schultz, Tatsuya Tsukahara, Sandra Martin Lorenzo, Valerie Nalesso, Alice H. Luo Clayton, Ted Abel, Jacqueline N. Crawley, Sandeep Robert Datta, Yann Herault

## Abstract

A microdeletion on human chromosome 16p11.2 is one of the most common copy number variants associated with autism spectrum disorder and other neurodevelopmental disabilities. Arbaclofen, a GABA(B) receptor agonist, is a component of racemic baclofen, which is FDA-approved for treating spasticity, and has been shown to alleviate behavioral phenotypes, including recognition memory deficits, in animal models of 16p11.2 deletion. Given the lack of reproducibility sometimes observed in mouse behavioral studies, we brought together a consortium of four laboratories to study the effects of arbaclofen on behavior in three different mouse lines with deletions in the mouse region syntenic to human 16p11.2 to test the robustness of these findings. Arbaclofen rescued cognitive deficits seen in two 16p11.2 deletion mouse lines in traditional recognition memory paradigms. Using an unsupervised machine-learning approach to analyze behavior, one lab found that arbaclofen also rescued differences in exploratory behavior in the open field in 16p11.2 deletion mice. Arbaclofen was not sedating and had modest off-target behavioral effects at the doses tested. Our studies show that arbaclofen consistently rescues behavioral phenotypes in 16p11.2 deletion mice, providing support for clinical trials of arbaclofen in humans with this deletion.

**One sentence summary:** Experiments across four laboratories found that arbaclofen rescued cognitive deficits in mouse models of 16p11.2 deletion, without sedation or significant off-target behavioral effects.

## Introduction

Human chromosome 16p11.2 microdeletion is one of the most common copy number variants associated with autism spectrum disorder (ASD) and other related neurodevelopmental disorders(*1*). Animal studies suggest that 16p11.2 deletion may share pathophysiology with Fragile X syndrome (FXS), namely similar behavioral phenotypes and similar changes in mGluR5-depedent synaptic plasticity and protein synthesis in the hippocampus(*2*). Treatments that improve symptomatology in FXS may therefore be of potential use for people with 16p11.2 deletion as well.

Data from a previous clinical trial suggest that the GABA-B receptor agonist, arbaclofen (R-baclofen), may improve symptomatology in some individuals with Fragile X syndrome and idiopathic autism, although the trial did not find any statistically significant differences on the primary endpoints(*3*). Arbaclofen is the pharmacologically-active enantiomer of racemic baclofen(*4*), showing greater potency than S-baclofen in a variety of biological and behavioral assays(*5*). Racemic baclofen is approved by both the FDA and the EMA (European Medicines Agency) for the treatment of spasticity associated with either multiple sclerosis or cerebral palsy and is commonly prescribed to children and adolescents with cerebral palsy(*6*). Arbaclofen appears to be quite safe, with sedation being the most common side effect(*7*).

A previous study in mouse models suggested that arbaclofen may rescue behavioral phenotypes in mice with a deletion on chromosome 7F3, the region syntenic to the human 16p11.2 microdeletion (referred to here as 16p11.2 del mice)(*8*). Three different mouse lines carrying highly overlapping but slightly different deletions in the syntenic region have been created; the three mouse lines are maintained on different genetic background strains (see Figure 1A) (*9–11*). In the Stoppel et al study, arbaclofen administration rescued object recognition memory and fear conditioning deficits in one of the lines of 16p11.2 del mice tested in the Bear laboratory, and rescued object location memory and social interaction deficits in the Crawley laboratory in a second, distinct mouse line(*8*). Taken together, these data suggest that arbaclofen may have efficacy in improving behavioral phenotypes in 16p11.2 deletion mice.

**Figure 1.**
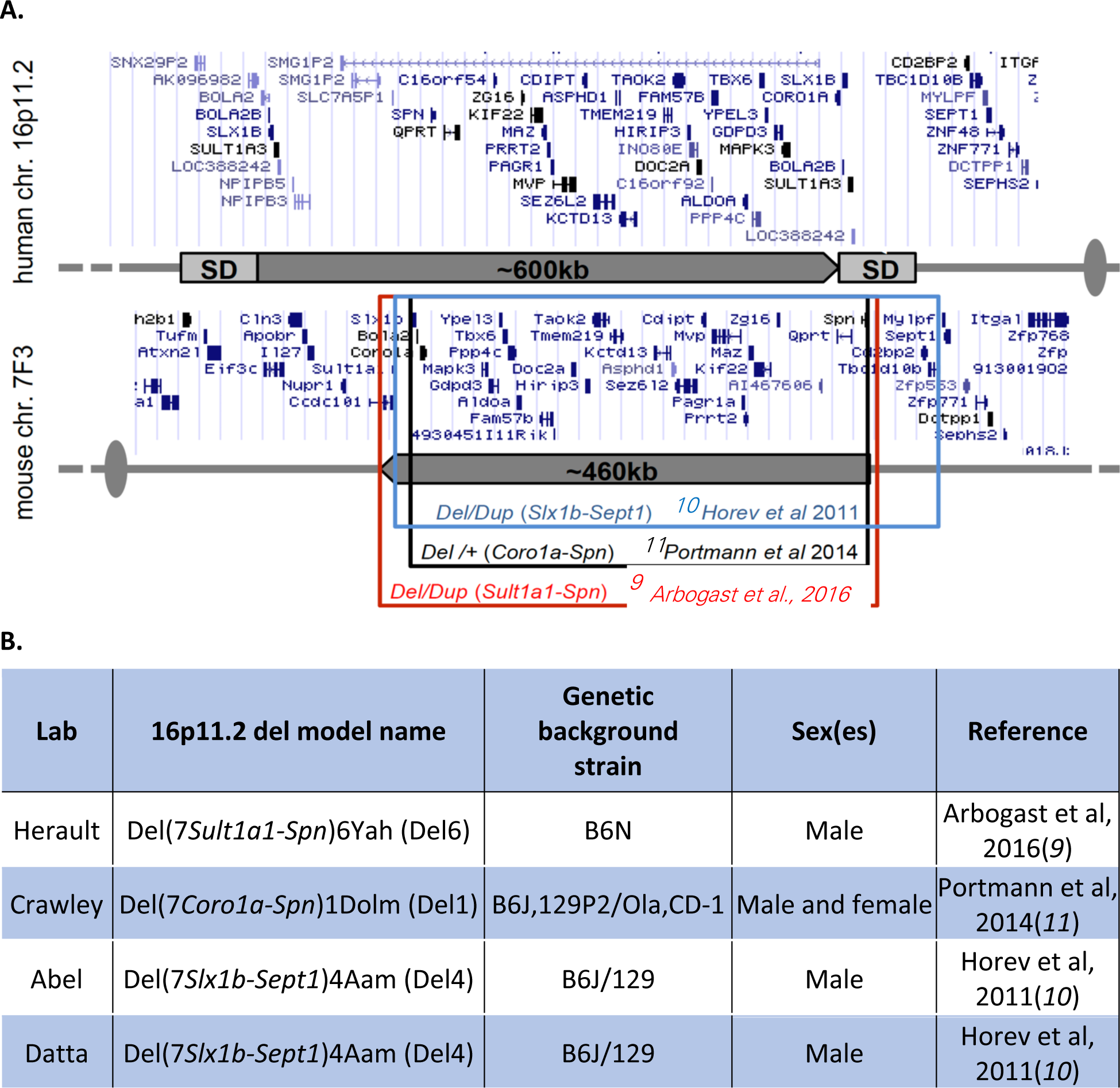
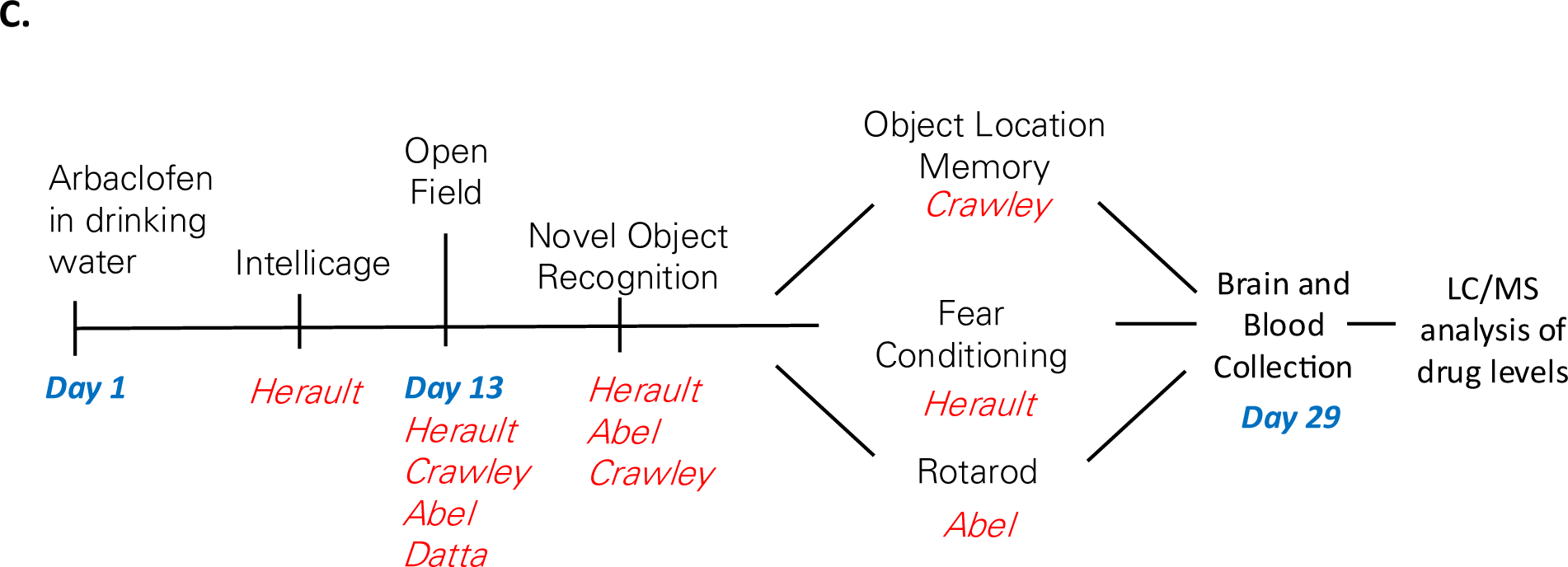
Study design. **A.** Schematic shows the locus of the 16p11.2 deletion in humans and the homologous 7F3 deletion in mice. Breakpoints of each of the 3 models of mice used in this study are indicated. Adapted from (*9*). **B.** Table listing the different 16p11.2 mouse models used in this study and the genetic background strains on which they were maintained. **C.** Experimental time model.

It is an unfortunately a reality in the field that mouse behavioral studies often produce inconsistent or irreproducible results, often attributed to inter-lab idiosyncrasies, differences in experimental or housing conditions, or differences in genetic background strain (*12, 13*). Even seemingly robust findings in rodent studies have failed to translate to successful clinical trials(*14, 15*). Concerted efforts to validate early scientific findings using rigorous methodological designs are crucial in addressing the recent paucity of success in translational research(*16*).

Therefore, we undertook a further study of arbaclofen’s effects on behavior in 16p11.2 del mice, taking advantage of the three different mouse lines available. In addition, we assembled a consortium of four labs to increase the robustness and reproducibility of our findings by using different points of analysis in different lab environments in which to observe the treatment effect.

## Results

To test the robustness and reproducibility of arbaclofen’s effects on behavior in 16p11.2 del mice, we assembled a consortium of 4 labs. Each lab worked with one of three mouse models of 16p11.2 deletion, Del1, Del4 and Del6, which were maintained on different background strains (Figure 1A). Adult mice (between 7-9 weeks old) were treated with one of three doses of arbaclofen in the drinking water (0.25, 0.5, or 1.0 mg/mL) for at least 12 days prior to behavioral testing. This dosing paradigm was based on results from a previous study of arbaclofen’s behavioral effects in 16p11.2 del mice (*8*). Previous studies of these mouse models of 16p11.2 deletion had described a variety of behavioral phenotypes, with variable results across models and across publications (*9–11, 17–19*). Based on these previous studies, the phenotype that seemed the most robust and reproducible in 16p11.2 del mice was a deficit in recognition memory (either in the novel object recognition, NOR, or object location memory, OLM, tasks), so we chose the NOR as our primary readout for examining the behavioral effects of arbaclofen. In addition to the NOR, each lab chose an additional behavioral test of interest that was administered following the NOR: OLM (Crawley), accelerating rotarod (Abel), or contextual fear conditioning (Herault) (Figure 1B). Because sedation is a potential side effect of arbaclofen, and would confound the interpretation of results from other behavioral tasks, we also included the open field test as a measure of general locomotion in the behavioral battery. The Datta lab looked solely at behavior in an open field, using a data-driven analysis of behavior(*20*).

In addition to the behavioral tasks, consumption of arbaclofen was measured (including in Intellicages by the Herault lab, see Methods), body weights were monitored throughout the study, and brain tissue was collected at the end of the study for measurement of arbaclofen content. Intellicage data show that mice of both genotypes, Del6 and wt, consumed less arbaclofen-containing water than those receiving plain water, an effect that appears to be dose-dependent (Figure S1A). This reduction in drinking normalized across days of treatment, with mice receiving the highest dose of arbaclofen indistinguishable from vehicle-treated mice by day 6 of treatment. A similar reduction in consumption of water in cages receiving arbaclofen was seen in both the Abel (Del4) and Crawley (Del1) labs (data not shown). The reduction in drinking observed in the Herault lab was accompanied by a slowing in the growth of body weights in arbaclofen-treated mice (Figure S1B). Again, similar trends in body weights were seen in the other three labs (data not shown). Note that mice of all three 16p11.2 deletion models generally had lower body weights than wildtype mice, as has previously been reported(*9, 11, 17*). HPLC-MS/MS analysis of brain tissue at the conclusion of the study revealed that arbaclofen was present at dose-dependent concentrations in the brains of treated mice (Figure S2).

16p11.2 deletion model mice have previously been reported to have impaired recognition memory in the NOR and OLM(*17, 18*), and arbaclofen has been reported to rescue these deficits(*8*). We found that two of the three mouse models of 16p11.2 deletion tested showed deficits in recognition memory in the NOR (Del 4 and Del6; Figure 2 A-C); in both, these deficits could be rescued by arbaclofen at all tested doses. Data from the Crawley lab did not show a deficit in recognition memory in the 16p11.2 Del1 deletion model mice in the NOR, nor in the OLM, in which they had previously been reported to show a deficit(*8, 18*). Raw exploration time data for both training and testing days of the task are shown in Figure S3.

**Figure 2.**
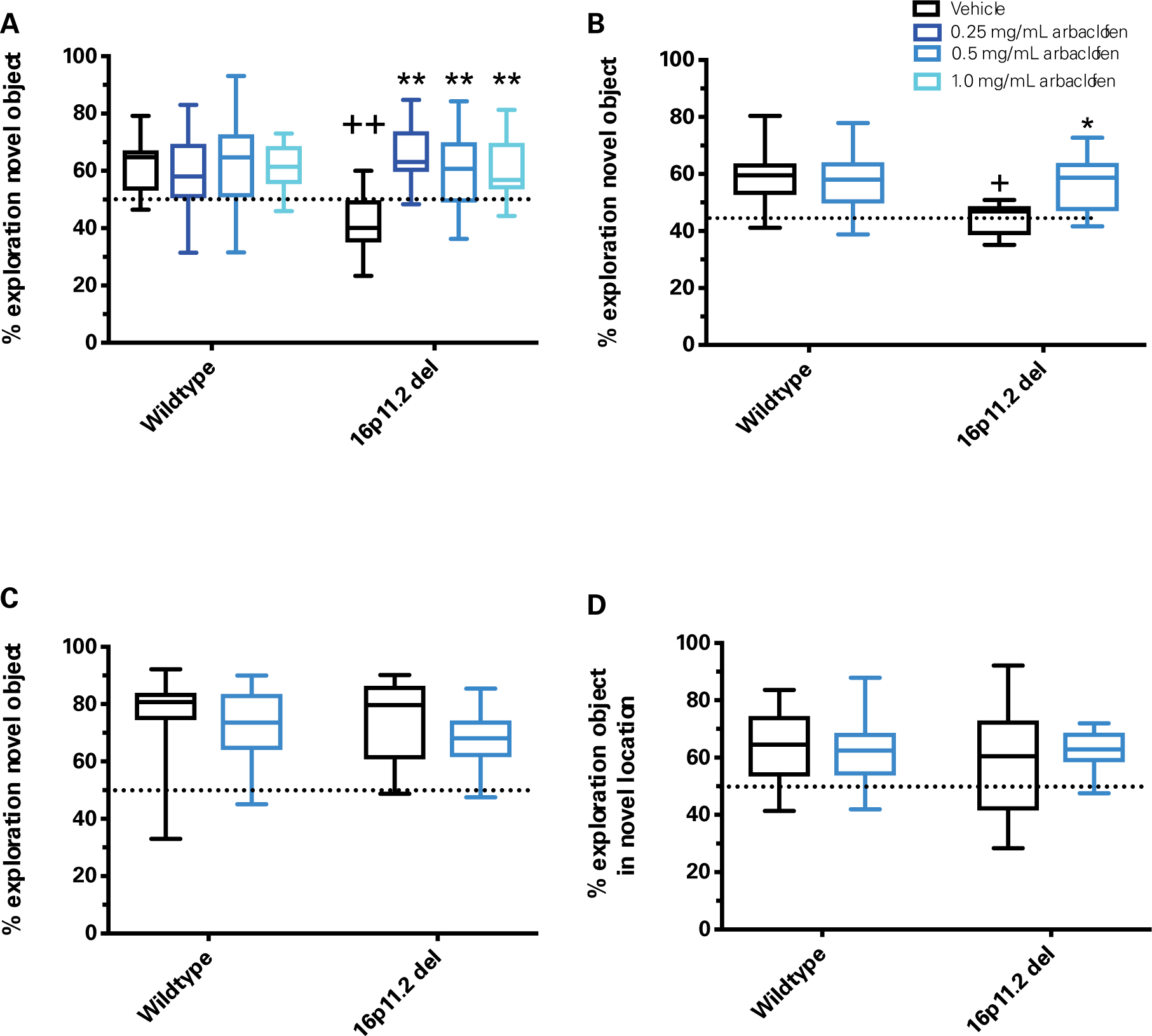
Arbaclofen rescues recognition memory deficits observed in two models of 16p11.2 deletion mice. Graphs are box and whisker plots; whiskers are maximum and minimum, boxes are interquartile range, and midline is median. **A-C.** During the training phase of the novel object recognition task, wt and 16p11.2 del model mice treated with vehicle or one of 3 doses of arbaclofen were allowed to interact with two identical objects. After a delay, mice were tested by exposure to one of the familiar objects and one novel object. Discrimination index (time spent exploring novel object/total object exploration time) for the testing phase is shown. Graphs are box and whisker plots; whiskers are maximum and minimum, boxes are interquartile range, and midline is median. Data were analyzed by 2-way ANOVA (genotype, treatment) with post-hoc t-tests (p values adjusted for multiple comparisons). **A:** Data collected in the Herault lab (3-hour delay between training and testing). Vehicle-treated wt mice show a significant preference for the novel object whereas vehicle-treated 16p11.2 Del6 deletion model mice show no significant preference for the novel object. Impaired novelty preference in 16p11.2 deletion model mice is rescued by all doses of arbaclofen. No main effect of genotype (F(1,95) – 2.513, *p*=0.1163), significant main effect of treatment (F(3,95)=4.048, *p*=0.0094), significant genotype x treatment interaction (F(3,95)=5.433, *p*=0.0017). Post-hoc t-tests (adjusted): wt/vehicle vs. 16p11.2 del/vehicle p=0.00014; 16p11.2 del/vehicle vs. 16p11.2 del 0.25 mg/mL p=0.0003; 16p11.2 del/vehicle vs. 16p11.2 del 0.5 mg/mL p= 0.0010; 16p11.2 del/vehicle vs. 16p11.2 del 1.0 mg/mL p= 0.0007. n= 14, 13, 13, 11, 13, 13, 14, 12. **B:** Data collected in the Abel lab (24-hour delay between training and testing). Vehicle-treated wt mice show a significant preference for the novel object whereas vehicle-treated 16p11.2 Del4 deletion model mice show no significant preference for the novel object. Impaired novelty preference in 16p11.2 Del4 deletion model mice is rescued by arbaclofen. Significant main effect of genotype (F(1,45)=6.625, *p*=0.0134), no main effect of treatment (F(1,45) = 3.239, *p*=0.0786), significant genotype x treatment interaction (F(1,45) = 5.890, *p*=0.0193). Post-hoc t-tests (adjusted): wt/vehicle vs. 16p11.2 del/vehicle p=0.00185; 16p11.2 del/vehicle vs. 16p11.2 del 0.5 mg/mL p= 0.0043. n = 16, 14, 7, 12. **C:** Data collected in the Crawley lab (1-hr delay between training and testing). Vehicle-treated wt and 16p11.2 Del1 deletion model mice each show significant preference for the novel object. No significant main effect of genotype (F(1,55) = 0.8994, p=0.3471), treatment (F(1.55) = 2.384, p= 0.1283) or genotype x treatment interaction (F(1,55) = 0.2346, p = 0.63). n = 18, 17, 14, 10. **D.** During the training phase of the object location memory task, wt and 16p11.2 Del1 del model mice treated with vehicle or a single dose of arbaclofen were allowed to interact with two identical objects. After a delay, mice were tested by exposure to the two trained objects, one in its previous location, and one moved to a novel location within the testing arena. Discrimination index (time spent exploring novel object/total object exploration time) for object location memory testing phase (1-hr delay between training and testing) for wildtype and 16p11.2 Del1 deletion model mice treated with vehicle or 0.5 mg/mL arbaclofen. Data collected in the Crawley lab. Vehicle-treated wt and 16p11.2 Del1 deletion model mice each show significant preference for the object in a novel location. Data were analyzed by 2-way ANOVA: no significant main effect of genotype (F(1,54) = 0.7187, p=0.4003, treatment (F(1,54)= 0.2433, p=0.6238) or genotype x treatment interaction (F(1,54)=0.5494, p=0.4618). n = 15, 17, 14, 12. + p < 0.01 vs. wt/vehicle, ++ p < 0.001 vs. wt/vehicle, * p < 0.01 vs. 16p11.2 del/vehicle, ** p < 0.001 vs. 16p11.2 del/vehicle.

The Datta lab used an unsupervised machine-learning approach to analyze behavior in an open field. This approach, MoSeq, uses 3D video imaging and applies a modified autoregressive-hidden-Markov model to parse behavior into sequences of sub-second motifs, or “syllables”(*20*). Comparing the frequency of each of 35 syllables across genotypes identified 7 syllables whose frequency significantly differed between wildtype and 16p11.2 deletion mice (Figure 3A). Treatment with the middle dose of arbaclofen (0.5 mg/mL) rescued the frequency of 6 of these 7 syllables in 16p11.2 deletion model mice, and the highest dose (1.0 mg/mL) rescued the frequency of all 7 (Figure 3B-H). The frequency with which 2-(bigrams, Supp Movies 1 and 2, Figure S4) and 3-syllable sequences (trigrams, Supp Movies 3 and 4, Figure S4) were expressed was also examined, revealing significant differences in the frequencies of 56 bigrams and 215 trigrams between wildtype and 16p11.2 deletion model mice. Arbaclofen treatment rescued differences in individual bigram and trigram frequency between wildtype and 16p11.2 deletion model mice (Figure 4A-F). A measure of total differences in the frequencies of bigrams (Figure 4E, Figure S5) or trigrams (Figure 4F, Figure S6) between genotypes also shows significant and dose-dependent rescue by arbaclofen.

**Figure 3.**
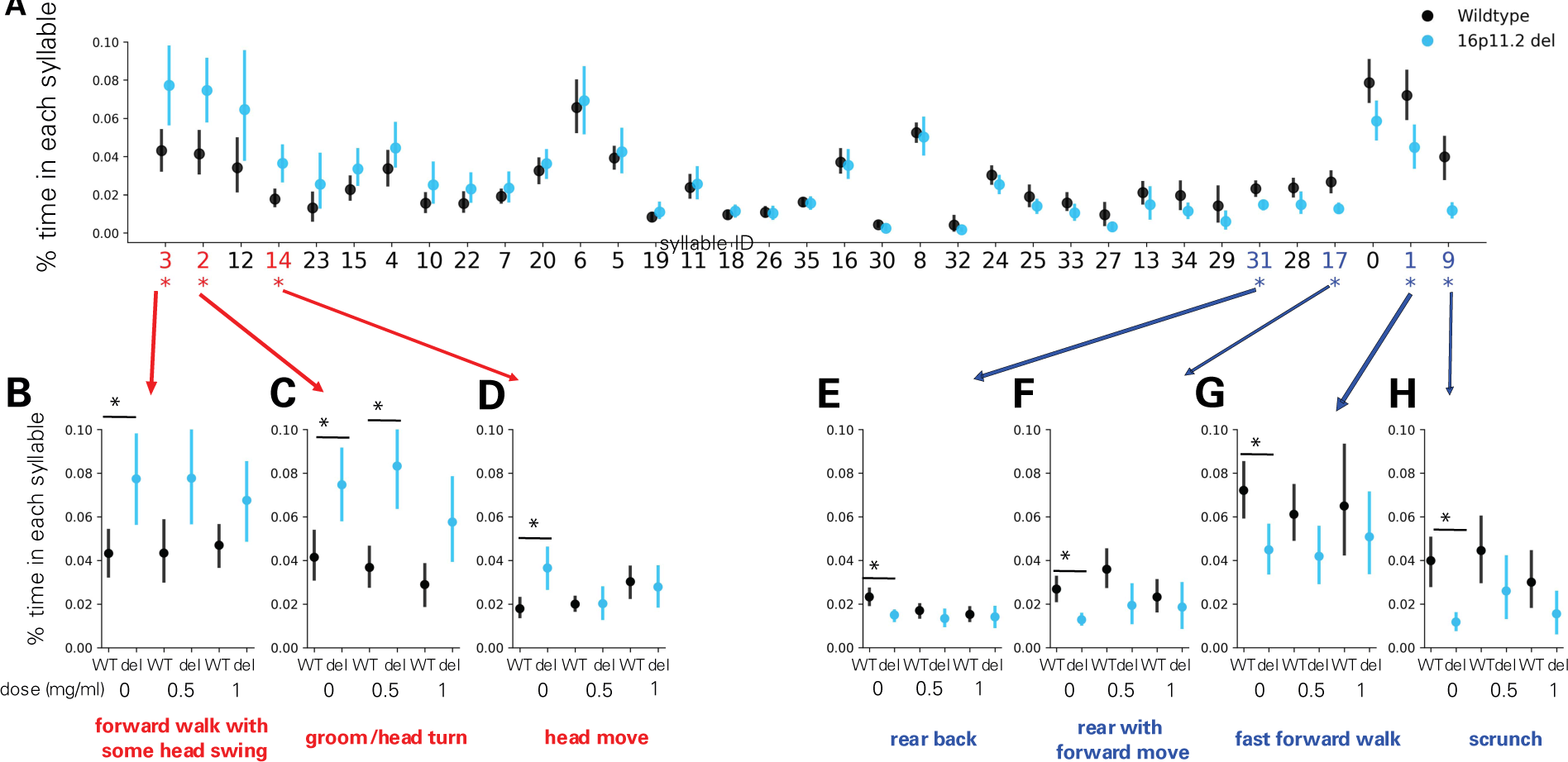
Differences in syllable frequency in 16p11.2 Del4 deletion model mice are rescued by arbaclofen treatment. **A-H.** Data collected in the Datta lab. n= 15 wt/vehicle, 12 16p del/vehicle, 27 wt/0.5 mg/mL arbac, 14 16p del/0.5 mg/mL arbac, 18 wt/1.0 mg/mL arbac, 11 16p del/1.0 mg/mL arbac**. A.** Ratio of the time in each syllable for vehicle-treated animals is plotted. Syllables are ordered by the magnitude of the difference between genotypes (largest differences between genotypes on the left and right ends). Circles indicate the population mean and error bars indicates 95% confidence interval estimated of 1000 bootstrapping. Asterisks indicate significantly different syllable (p< 0.05 with Z-test in which standard deviation was estimated by 1000 bootstrapping and Benjamini-Hochberg FDR correction was performed). Red numbers indicate syllables that are significantly up-regulated in 16p11.2 Del4 deletion model mice and blue numbers indicate syllables that are significantly down-regulated in 16p11.2 Del4 deletion model mice. **B-H.** Ratio of the time spent performing each of the syllables significantly different between 16p11.2 Del4 deletion model mice and wildtype treated with one of 3 doses of R-baclofen (0 (vehicle), 0.5, 1.0 mg/ml). Annotations indicate human descriptions of the behavior contained in each syllable. Asterisks indicate significant difference (analysis as in A). Circles indicate the population mean and error bars indicates 95% confidence interval estimated of 1000 bootstrapping.

**Figure 4.**
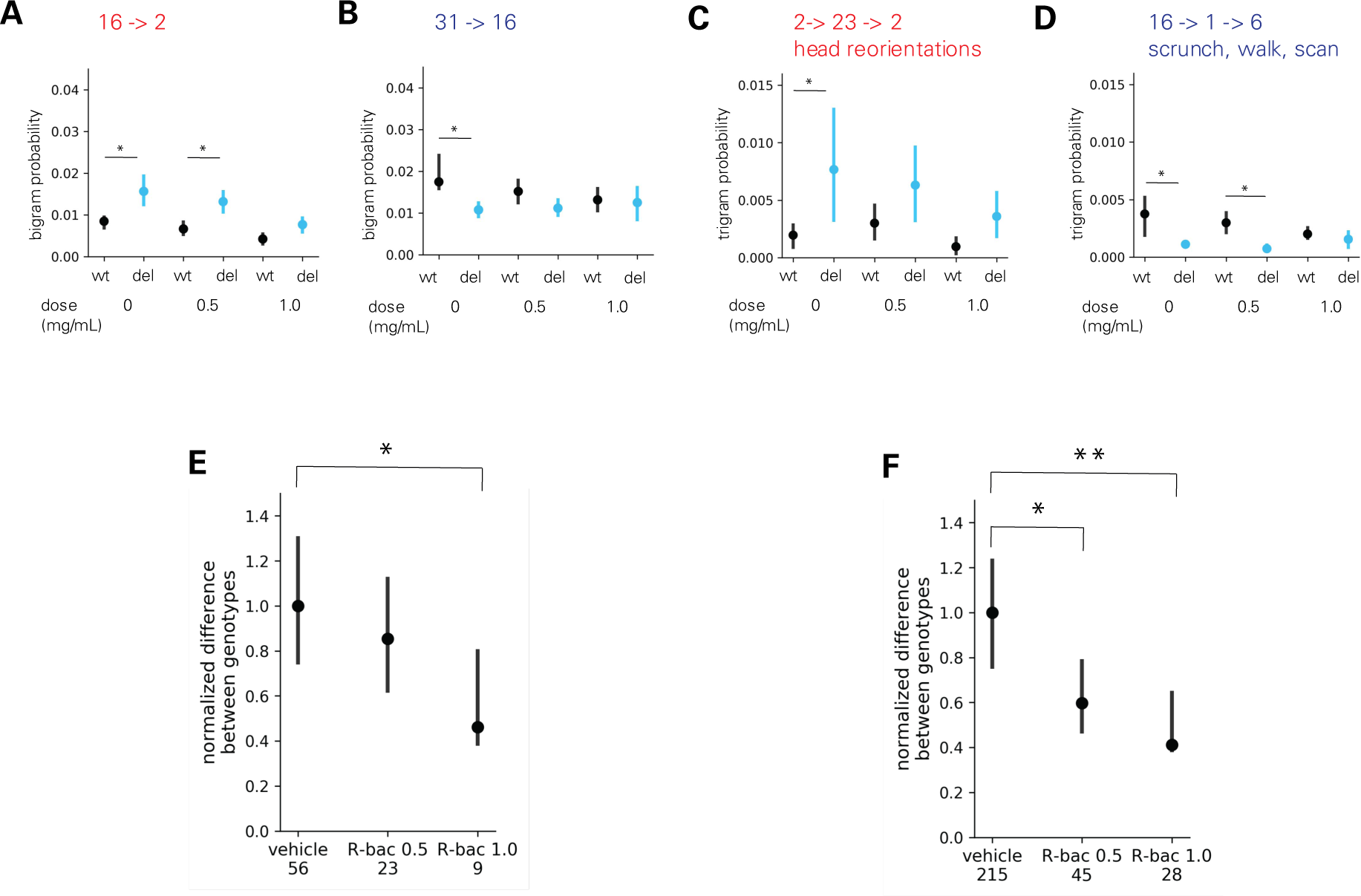
Differences in transition probability in 16p11.2 Del4 deletion model mice are rescued by arbaclofen treatment. **A-F.** Data collected in the Datta lab. n= 15 wt/vehicle, 12 16p del/vehicle, 27 wt/0.5 mg/mL arbac, 14 16p del/0.5 mg/mL arbac, 18 wt/1.0 mg/mL arbac, 11 16p del/1.0 mg/mL arbac **A-B.** Examples graphs showing probabilities of expressing individual bigrams across genotype and treatment groups. Error bars indicate 95% confidence intervals from 1000 bootstrapping and asterisks indicate non-overlap of confidence intervals between Wildtype and 16p11.2 Del4 del for each condition. **C-D.** Examples graphs showing probabilities of expressing individual trigrams across genotype and treatment groups. Error bars indicate 95% confidence intervals from 1000 bootstrapping and asterisks indicate non-overlap of confidence intervals between wildtype and Del4 16p11.2 deletion model mice for each condition. **E-F.** Summed differences in probabilities of 56 bigrams (E) or 215 trigrams (F) differentially used between wildtype and Del4 16p11.2 deletion model mice in each condition. Note that all data is scaled to the total differences observed between wildtype and Del4 16p11.2 deletion model mice treated with vehicle (0 mg/ml). Bigrams/trigrams detected more than chance were identified by Monte Carlo randomization for each condition (50000 times: p < 5e-5 for bigrams. 100000 times: p<2e-5 for trigrams, one sided). The union of such bigrams/trigrams in different groups was then compared by bootstrap test (1000 times). Significantly different bigrams/trigrams were identified if 95% confidence intervals were not overlapped. Number of bi/trigrams that are still significant in R-baclofen groups out of 56/215 differential ones in vehicle condition are listed underneath x-axis. P-values for the summed difference were calculated as follows: p = (# of bootstraps where the value for group A is less than that for group B + 1) / (# of bootstraps). Group A = vehicle, B = R-bac 0.5 or R-bac 1.0. p values were subjected to Bonferonni correction, and corrected p values <0.05 were considered significant. *p < 0.05, **p<0.01 between vehicle and drug conditions. Error bars indicate 95% confidence interval from 1000 bootstraps. Number of bi/trigrams that are still significant in R-baclofen groups out of 56/215 differential ones in vehicle condition are listed underneath x-axis.

Previous studies have reported deficits in contextual fear conditioning in the Del1 model of 16p11.2 deletion mice(*8*). The Herault lab tested whether any deficits in contextual fear conditioning in the 16p11.2 Del6 deletion model mice could be rescued by arbaclofen. Wildtype and 16p11.2 deletion model mice were trained to associate a foot shock with one context, and then tested the following day in either the trained context or a novel context (see methods). There was no significant difference in freezing behavior in 16p11.2 Del6 deletion model mice compared to wildtype mice (Figure 5). Arbaclofen also did not significantly alter freezing behavior in either genotype.

**Figure 5.**
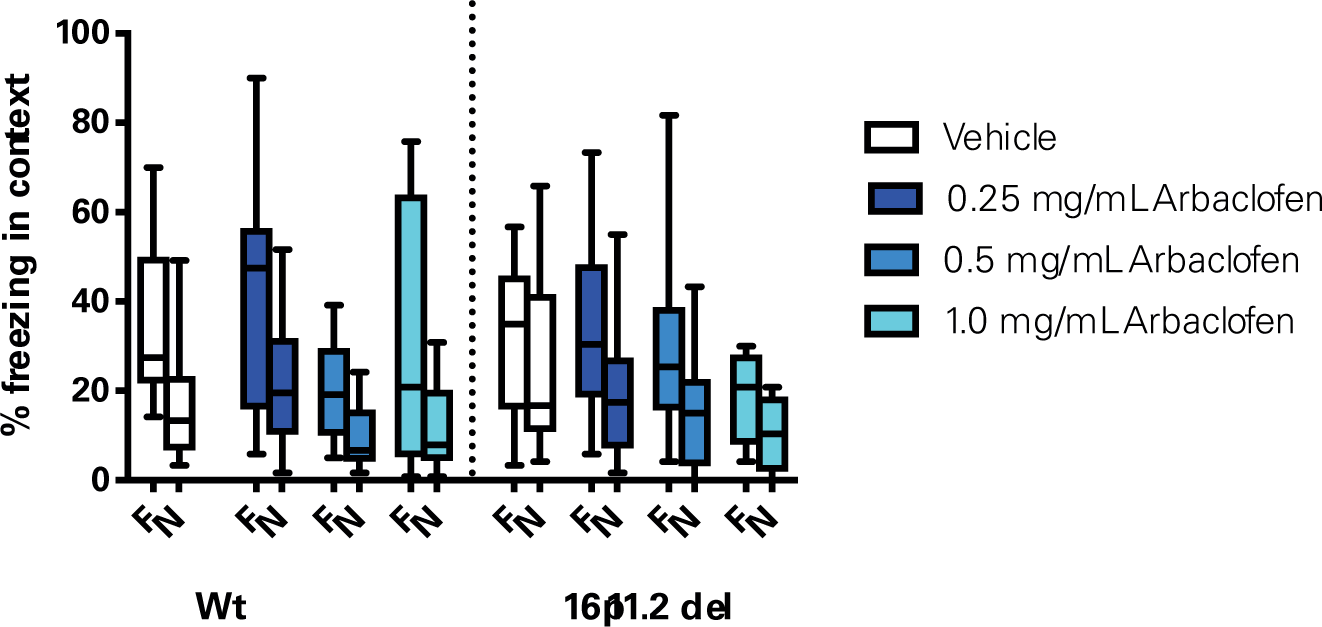
No difference in contextual fear conditioning in 16p11.2 Del6 deletion model mice. Wt and 16p11.2 Del6 deletion model mice treated with one of 3 doses of arbaclofen were exposed to one context in which they were given a single foot shock. The following day, mice were exposed to the training context (familiar, F) and to a novel context (N) in counterbalanced order, without receiving a foot shock, for 8 minutes. Percent time spent freezing in each context is shown. Data were collected in the Herault lab. Graphs are box and whisker plots; whiskers are maximum and minimum, boxes are interquartile range, and midline is median. Data were analyzed with 3-way RMANOVA (genotype and treatment as between-subjects variables, context as within-subject variable): significant main effect of context (F(1,104) = 81.275, *p*<0.0001), no significant main effect of genotype (F(1,104) = 0.704, p=.4034), treatment (F(3,104) = 1.607, p=.1923), or significant genotype x treatment (F(3,104) = 1.168, p= 0.3257) or genotype x treatment x context interaction (F(3, 104) = 1.280, p =0.2851). F = Familiar context; N = Novel context. n = 15, 14, 13, 10, 15, 14, 14, 12.

Enhanced learning in a more challenging version of the accelerating rotarod task has also been previously reported in 16p11.2 deletion model mice(*17*) as well other mouse models of autism (*21–26*). The Abel lab tested whether this learning phenotype in 16p11.2 Del4 deletion model mice was altered by treatment with arbaclofen. Wildtype and 16p11.2 deletion model mice were trained on the accelerating rotarod at low speeds (4-40 rpm/5 min) for 3 trials on day 1 (standard paradigm), and then at higher speeds (8-80 rpm/5 min) for 3 trials on day 2 (more challenging paradigm). As was previously reported (*17*), 16p11.2 Del4 deletion model mice had significantly longer latencies to fall on both days (Figure 6). However, arbaclofen did not affect rotarod performance in either wildtype or 16p11.2 deletion model mice.

**Figure 6.**
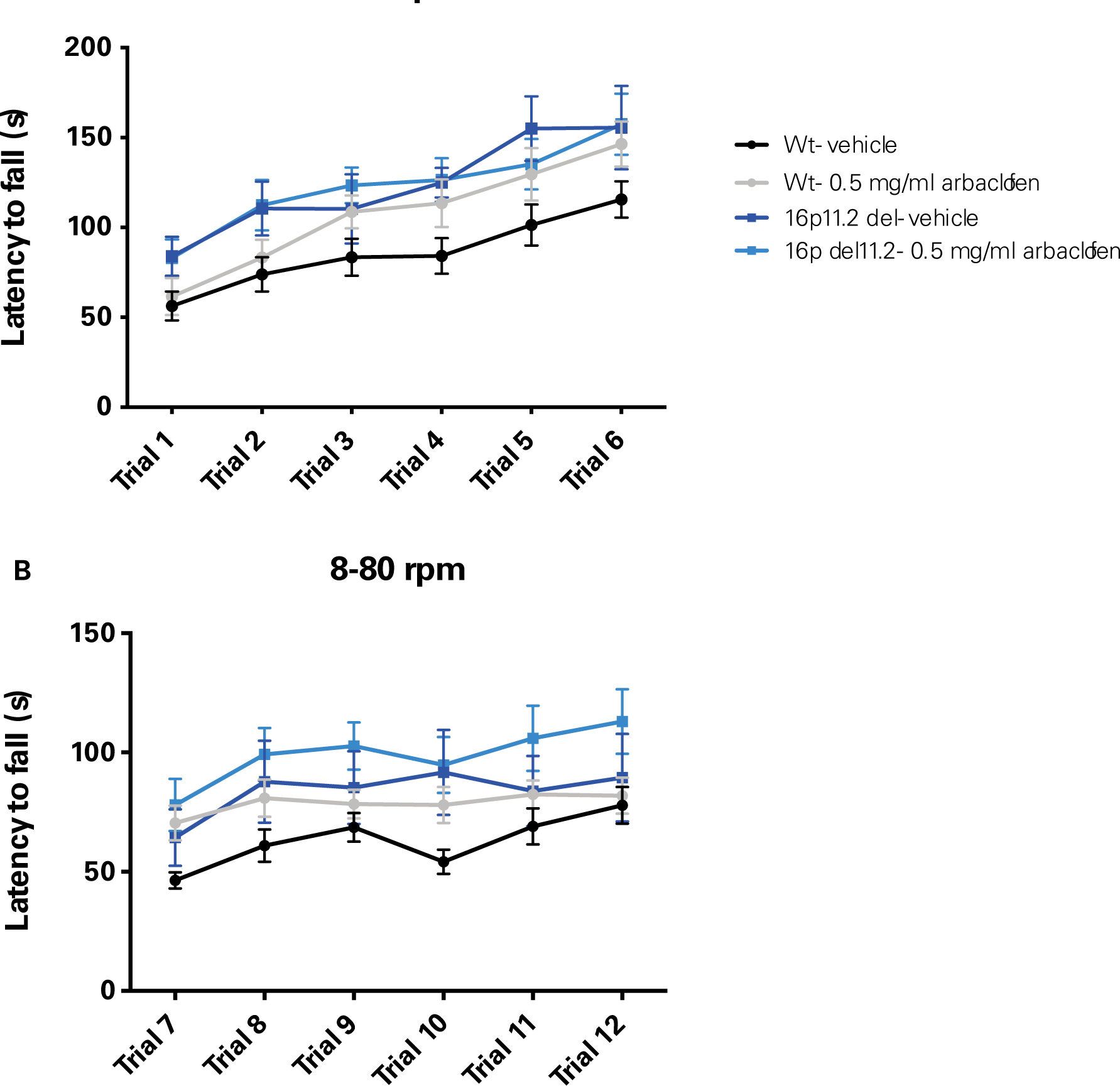
16p11.2 Del4 deletion model mice show enhanced learning in the accelerated rotarod that is unaffected by arbaclofen. Wildtype and 16p11.2 Del4 deletion model mice treated with a single dose of arbaclofen were trained on the accelerating rotarod at low speeds (4-40 rpm/5 min) for 3 trials on day 1, and then at higher speeds (8-80 rpm/5 min) for 3 trials on day 2. Latency to fall is shown for day 1, 4-40 rpm (A) and day 2, 8-80 rpm (B). Data were collected in the Abel lab. Points are means with error bars indicating SEM. Data were analyzed with 3-way RMANOVA (genotype and treatment as between-subjects variables, trial as within-subject variable): A. 3-way RMANOVA: significant effect of trial (F(5,48)=51.6, p<0.001) and genotype (F(1,48)=5.82, p<0.05) but not treatment (F(1,48)=0.89, p=0.35) or genotype x treatment interaction (F(1,48)=0.96, p=0.33). **B**. 2-way RMANOVA: significant main effect of trial (F(5,48)=14.0, p<0.001) and genotype (F(1,48)=5.54, p<0.05) but not treatment (F(1,48)=3.15, p=0.08) or genotype x treatment interaction (F(1,48)=0.001, p=0.97). n= 17, 15, 8, 12.

As a GABA(B) agonist, arbaclofen might produce sedation. All four labs examined open field behavior to assess potential effects of arbaclofen on general locomotor behavior. The data were inconsistent across labs, but overall do not show a sedative effect of arbaclofen at the doses that rescued other behavioral phenotypes (Figure 7). In the Herault lab, 16p11.2 Del6 deletion model mice showed increased locomotor activity overall, as has previously been described(*8–11*), and there was a significant effect of arbaclofen to increase activity (Figure 7A). In the Abel lab, there were no significant differences across genotypes (Del4 or wt) or treatment groups (Figure 7B). In the Crawley lab, there was a significant decrease in activity in the 16p11.2 Del1 deletion model mice, but no effect of treatment (Figure 7C). Analysis of rearing behavior and time spent in the center of the open field reveal equally inconsistent results across the three models (Figure S7 and S8). Analysis of mouse velocity and overall distance traveled from the Datta lab also does not show significant sedative effects of arbaclofen, whereas risperidone, a drug prescribed to treat irritability in ASD, does significantly decrease velocity and distance traveled in the 16p11.2 deletion model mice (Figure S9).

**Figure 7.**
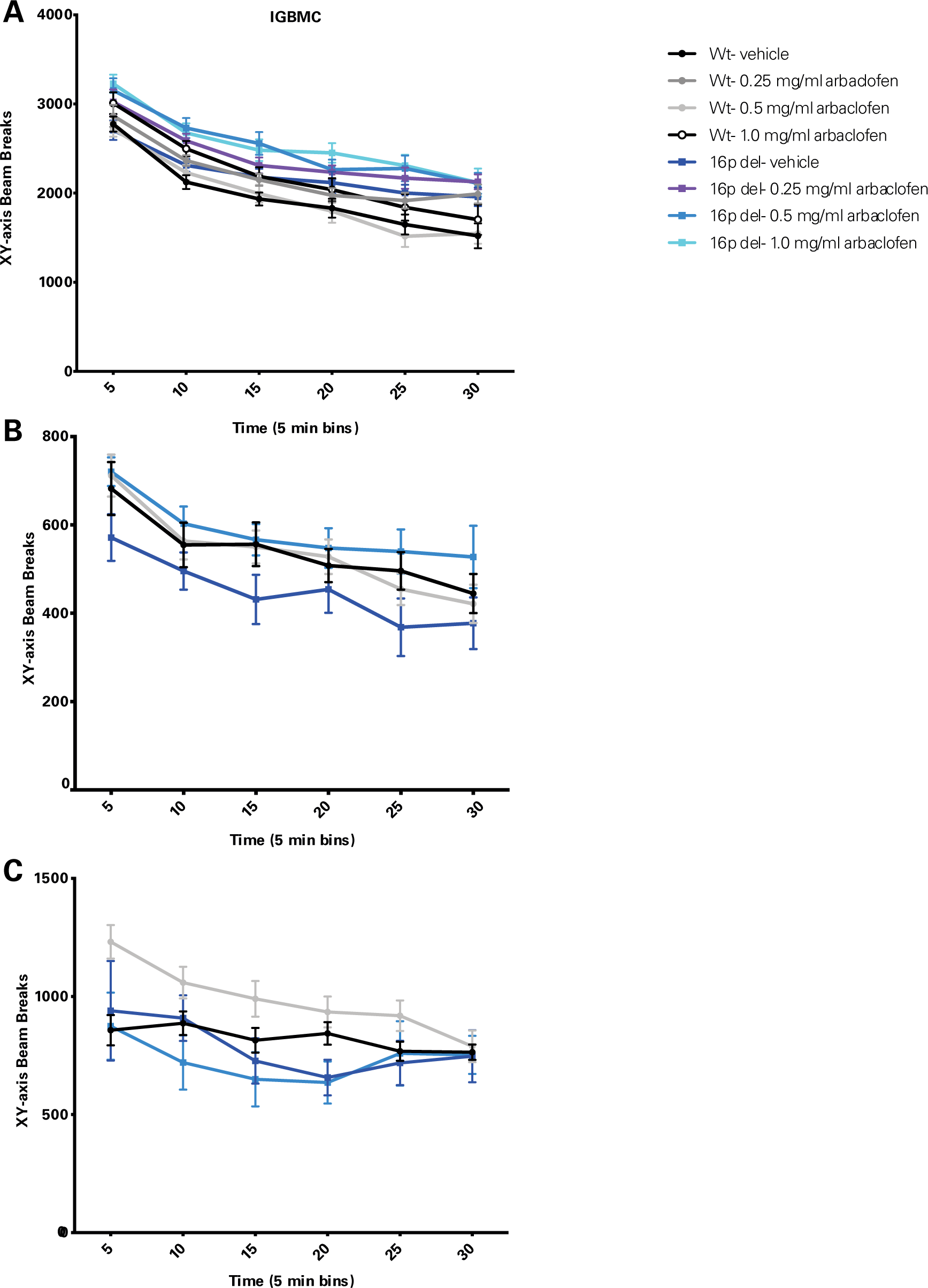
Arbaclofen does not reduce locomotor activity. Wildtype and 16p11.2 deletion model mice treated with one of 3 doses of arbaclofen were exposed to an open field for 30 min. Activity counts (measured by infrared beam breaks in the x and y planes) are plotted in 5-minute bins. Points are means with error bars indicating SEM. Data were analyzed with 3-way RMANOVA (genotype and treatment as between-subjects variables, time as within-subject variable): **A.** Data were collected in the Herault lab (Del6 model). 3-way RMANOVA: significant main effect of time (F(5,101) = 200.958, p<.0001), significant main effect of genotype (F(1,101) = 6.056, p=0.0156), significant main effect of treatment (F(3, 101) = 3.799, p=0.0126), significant time x genotype interaction (F(5, 101) = 3.465, p=0.0043). n = 14, 14, 14, 11, 14, 14, 15, 12. **B.** Data were collected in the Abel lab (Del4 model). 3-way RMANOVA: significant main effect of time (F(5, 46) = 34.126, p=2×10-^16^); no significant main effects of genotype (F(1,46) = 0.030, p=0.862), treatment (F(1,46) = 1.215, p = 0.276), or significant interactions. n = 17, 15, 7, 11. **C.** Data were collected at in the Crawley lab (Del1 model). 3-way RMANOVA: significant main effect of time (F(5, 62) = 7.699, p = 7.69 × 10^-7^); significant main effect of genotype (F(1, 62) = 5.358, p = 0.024); no main effect of treatment (F(1, 62) = 1.466, p = 0.2305) or significant interactions. n= 19, 20, 14, 13.

In addition to sedation, we were interested in other potential off-target behavioral effects induced by arbaclofen treatment. To address this question, we examined the effects of arbaclofen and risperidone on behavioral transitions between pairs of syllables (bigrams and trigrams) using MoSeq in wildtype mice in the Datta lab (Figure 8). Arbaclofen did have dose-dependent effects on behavioral transitions in wildtype mice (Figure 8A), but both doses of arbaclofen affected syllable sequences significantly less than did risperidone (Figure 8B-C).

**Figure 8.**
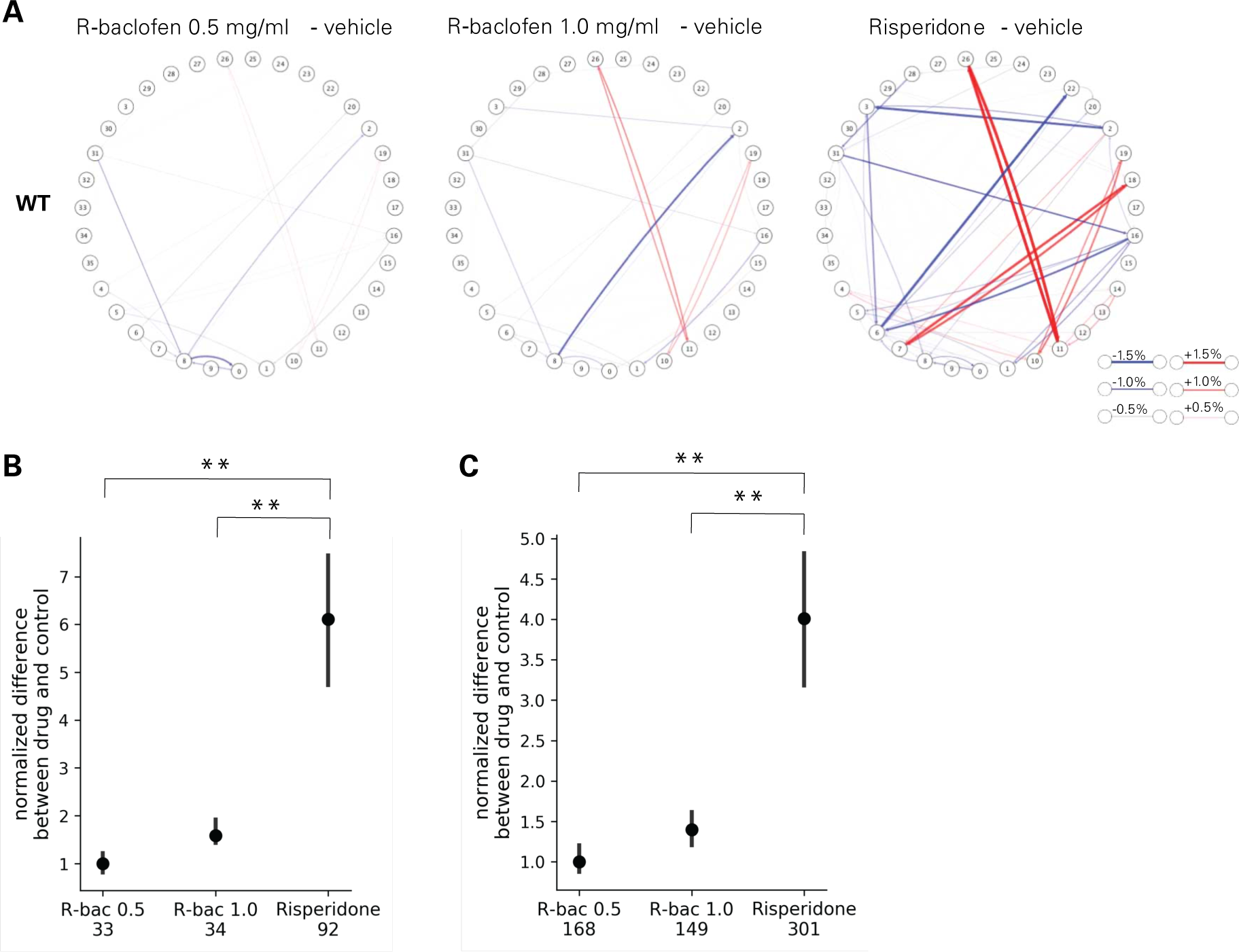
Compared to risperidone, arbaclofen does not produce major off-target behavioral effects. **A-C.** Data were collected in the Datta lab (Del4 model). **A.** Plots show the difference in bigram probabilities between drug and control conditions in wildtype mice. Each node represents a syllable (as numbered) and each line indicates a transition from one syllable to another. Line thickness indicates the magnitude of difference between treatments. Red lines indicate bigrams up-regulated with drug treatment and blue lines indicate bigrams down-regulated by drug treatment. n= 15 wt/vehicle; 27 wt/0.5 mg/kg arbaclofen; 18 wt/1.0 mg/kg arbaclofen; 6 wt/saline; 13 wt/risperidone **B-C.** Summed differences in probabilities of all differentially-used bigrams (B) or trigrams (C) in wildtype mice, comparing drug-treated (0.5 or 1.0 mg/mL arbaclofen or risperidone) and vehicle-treated mice. Bigrams/trigrams detected more than chance were identified by Monte Carlo randomization for each condition (50000 times: p < 5e-5 for bigrams. 100000 times: p<2e-5 for trigrams, one sided). The union of such bigrams/trigrams in different groups was then compared by bootstrap test (1000 times). Significantly different bigrams/trigrams were identified if 95% confidence intervals were not overlapped. Number of bi/trigrams that are significantly different between drug and vehicle are listed underneath x-axis. P-values for the summed difference were calculated as follows: p = (# of bootstraps where the value for group A is less than that for group B + 1) / (# of bootstraps). Group A = Risperidone, B = arbaclofen 0.5 mg/mL or arbaclofen 1.0 mg/mL. p values were subjected to Bonferonni correction, and corrected p values <0.05 were considered significant. **p<0.01 between Risperidone and R-baclofen conditions. n= 15 wt/vehicle; 27 wt/0.5 mg/kg arbaclofen; 18 wt/1.0 mg/kg arbaclofen; 6 wt/saline; 13 wt/risperidone

## Discussion

We rigorously assessed the efficacy of arbaclofen to rescue behavioral phenotypes in 16p11.2 deletion mice across four labs using one of three different mouse models of the deletion, with different genetic constructs and maintained on different background strains (Figure 1A). We found that, in two of three labs using traditional behavioral paradigms, arbaclofen robustly rescued recognition memory deficits at all tested doses (Figure 2), replicating published results from an independent lab(*8*). The fourth lab used a data-driven approach, which parses behavior at sub-second resolution, showing that arbaclofen rescued differences in exploratory behavior in the open field (Figures 3 & 4). 16p11.2 Del6 deletion mice did not show a deficit in contextual fear conditioning (Figure 5), contrary to previous findings in the Del1 model (*8*). Nevertheless, 16p11.2 Del4 deletion mice did show improved learning in a challenging accelerated rotarod paradigm (Figure 6), which was not rescued by arbaclofen. In control experiments, arbaclofen did not seem to be sedating (Figure 7) and had modest off-target behavioral effects at tested doses (Figure 8). Overall, arbaclofen shows consistent behavioral rescue under testing strategies spanning multiple laboratories and mouse lines, as well as orthogonal analytical approaches.

Our consortium offers an example of how to develop and execute a rigorous test of preclinical efficacy of a potential pharmacological therapy. Mouse behavioral testing is subject to many potential confounds and limitations and results of behavioral experiments in one lab or one strain of mice are often hard to replicate in other labs or other background strains (*12, 13*). Our consortium sought to avoid some of these pitfalls by testing arbaclofen’s effects across multiple labs and using multiple mouse models with different genetic constructs and on different background strains(*9–11*). In addition, given the highly constrained nature of conventional (construct-based) mouse behavioral paradigms, some effects of a drug (both intended and unintended) may be missed. Our consortium therefore included both conventional and novel data-driven approaches(*20*) to measure mouse behavioral changes in response to arbaclofen.

When studies fail to see effects of a drug on behavior, one potential explanation is often that the drug has not reached its site of action (e.g. did not cross the blood brain barrier). In this study, HPLC-MS/MS analysis of brain tissue at the conclusion of the study showed that arbaclofen had indeed crossed the blood brain barrier and was present at dose-dependent concentrations in the brains of treated mice (Figure S2). This finding that oral treatment with arbaclofen does lead to presence of the drug in the brain is in concordance with previous findings(*5*). We did see some quantitative differences in the arbaclofen levels in the brains of treated mice across the four labs. These differences are likely due to variability inherent in the passive administration procedure (drug present in the drinking water) we used, or other procedural differences between labs, such as the exact timing of brain collection relative to when mice were removed from their home cages (and therefore, their arbaclofen access ended).

Arbaclofen rescued behavioral phenotypes in two 16p11.2 del models tested, Del4 and Del6, in the NOR (Figure 2). In the third lab (Crawley), 16p11.2 Del1 mice did not show a baseline phenotype in this task following vehicle treatment. They also did not show a baseline phenotype in the related OLM task, contrary to previous reports(*8, 18*). While the difference in behavioral phenotypes seen in the mutant mice could certainly be attributed to differences in genetic constructs and/or background strain of the mouse models, the Crawley lab also used both male and female mice, whereas the Abel and Herault labs used only male mice. In the NOR, there is an additional potential explanation for the differences seen: the delay period between training and testing sessions also differed across labs (Herault: 3 hr, Abel: 24 hr, Crawley: 1 hr). Differences in overall object exploration times seen across labs are also likely attributable to differences in mouse models/background strains, test conditions, and for the testing phase, delay period.

Although arbaclofen rescued deficits in the NOR in the 16p11.2 del mice in two of three studies, it did not reverse the additional phenotype observed in the Abel lab, in which mutant mice showed improved performance on a more challenging version of the accelerating rotarod paradigm (Figure 6). The lack of rescue in this case could be due to a difference in brain regions mediating these behaviors, with the NOR thought to depend on hippocampal function(*27*) whereas the rotarod is thought to reflect striatal function(*28, 29*). These tasks also tap into different constructs, with the NOR assessing cognitive function and the rotarod assessing motor function. It is also worth pointing out that the rotarod phenotype in the 16p11.2 del mice is an improvement, rather than a deficit, in performance, and thus it is possible that the lack of effect of arbaclofen could reflect some degree of ceiling effect.

The 16p11.2 del mice have previously been reported to be hyperactive(*8–11*). We found inconsistent results in our studies (Figure 7). In the Herault lab’s experiments, there was a significant main effect of genotype on horizontal activity in the open field, with Del6 mutant mice showing higher activity than wildtype mice, consistent with previous findings. However, in the Crawley lab, Del1 mutant mice showed lower overall horizontal activity in the open field, and in the Abel lab there was no significant effect of Del4 genotype. We also studied the effects of arbaclofen treatment on activity levels given concerns that arbaclofen, as a GABA agonist, might be sedating. Again, we found inconsistent results across labs, but if anything, arbaclofen treatment seemed to have a slight activating effect, with significant effects of treatment seen on horizontal activity in the Herault lab, and on vertical activity (rearing) in the Abel lab (Figure 7). These inconsistent results do not justify a strong conclusion regarding arbaclofen’s activating effects, but they do seem to rule out confounding sedative effects of the drug at the doses employed.

Overall, our findings show rigorous and reproducible effects of arbaclofen to rescue behavioral phenotypes in 16p11.2 del model mice across many different experimental and biological conditions. These results lend support to the potential efficacy of arbaclofen in humans with 16p11.2 deletion. Indeed, Clinical Research Associates is now conducting a clinical trial of arbaclofen in this population (registered at <u>clinicaltrials.gov</u> as NCT04271332), informed by this preclinical study. We hope future such forward- and back-translation between animal models and human studies will continue to advance the field towards effective interventions in this and other neurodevelopmental disorders.

## Materials and Methods

### Herault Lab (IGBMC)

#### Ethical statement

Experimental procedures were approved by the French Ministère de l’Enseignement Supérieur, de la Recherche et de l’Innovation under the accreditation number APAFIS#9290-2017031617456047v4.

#### Mice and housing conditions

The B6N Del(7*Sult1a1-Spn*)6Yah, referred here as Del6Yah/+, as previously described in (*9*), was maintained on a pure C57BL/6N genetic background. The cohorts of B6NDel6Yah/+ mice were collected after four rounds of in vivo fertilization to obtain up to 15 males in four groups with Del/+ and wt littermates. The production was done to assure a low heterogeneity in age. Only male mice were used in this study. At four weeks of age, male mice of both genotypes were weaned from several litters and placed into groups of 8-15 individuals and housed in large cages of 1400 cm^2^ (Polycarbonate type IVS, Genestil), where they had free access to water and diet (D04 chow diet, Safe, Augy, France). At the age of four weeks, they were transferred from the animal facility to the phenotyping area. Animal bedding (AB3, ANIBED, Pontvallain, France) was changed once a week. The temperature was maintained at 21±2 °C, with a 12 hr light/12 hr dark cycle (lights on at 07:00hr). At week 10, an RFID transponder (PeddyMark Ltd; type of transponder: DataMars (12mm x 2mm); ISO11784/1175) was implanted by subcutaneous injection into the dorso-cervical region of mice under isoflurane inhalation anesthesia. Three days after RFID implantation, mice were housed in the Intellicage monitoring system (TSE Systems GmbH, Bad Homburg, Germany) for 14 days in which they received either water or one of multiple doses of arbaclofen (see below). The day before the first day of behavioral testing, the animals were transferred in groups of four (all receiving the same arbaclofen dose) to 39 x 20 x16 cm cages (Green Line, Techniplast, Italy). The body weights of animals were recorded once per week (the same day of the week at the same time) from 4-13 weeks of age.

#### Drug Treatment

Arbaclofen (R-baclofen) was provided as a generous gift from Clinical Research Associates. Arbaclofen was administered *ad lib* in home cage drinking water at a concentration of 0.25, 0.5, or 1.0 mg/ml to starting at day 2 of Intellicage housing (10 weeks of age) and lasting throughout behavioral testing to the end of the study (29 days total). Water bottles were changed daily.

#### Behavioral testing

Mice underwent behavioral testing in the following order: open field (day 13, 14, 15, 16, or 17 of arbaclofen treatment)), novel object recognition task with a three hr retention delay (test day on day 19, 20, 21, 22, 23, or 24) and fear conditioning (test day on day 27, 28, 29, or 30). Testing of each cohort of mice was distributed across multiple days for each test, with at least one day between behavioral tests for a given mouse.

Behavioral experiments were conducted between 10 and 14 weeks of age. Behavioral testing was performed between 08:00 and 16:00hr. On testing days, animals were placed in the experimental room antechambers 30 min before the start of the experiment. The order of testing the mice was randomized by cage (order of cages randomized, order of mice within each cage randomized). The investigator performing behavioral tests was blind to the genotype and treatment of each mouse. Mice were excluded from behavioral testing and analysis if they were wounded or deceased (1 wt,0.25 mg/mL; 3 wt,0.5 mg/mL; 3 wt,1.0mg/mL; 1 del,0.25 mg/mL; 4 del,0.5 mg/mL; 2 del 1.0mg/mL) during the course of the study.

#### Intellicage system

Each IntelliCage (IC) (58cm x 40cm x 20.5cm) contained in each corner a triangular recording chamber (15cm x 15 cm x21cm) accessible through an open doorway, which recorded the identity of the radio-frequency identification (RFID) tag of the mouse passing through it. Each chamber contained two openings of 13 mm diameter, giving access to a water source. For this experiment, one drinking bottle was placed in one of the chambers, with access controlled by motorized doors. Animals could gain access to water via a nose-poke, which was recorded by photobeams. The number and duration of tongue-contacts on each drinking bottle were recorded by a lickometer. IC apparatuses were used simultaneously in the experiment for each group of treatment with 8-12 animals (mixed genotype) in each IC. During the first two days, animals were allowed to habituate to the IC and to localize the water source. During the next 12 days, arbaclofen or vehicle treatment was given in drinking water at a concentration of 0.25/0.5/1 mg/ml. The number of chamber visits and nose-pokes were measured to estimate spontaneous activity, while number of licks were used to measure the consumption of arbaclofen in drinking water during the 12-day period. The data from the IC system were analyzed using software (New Behavior AG, Zurich, Switzerland, www.newbehavior.com) to determine spontaneous activity and drinking behavior in group-housed mice.

#### Open field

Mice were tested in automated open fields (44.3 x 44.3 x 16.8 cm) made of PVC with transparent walls and a black floor, and covered with translucent PVC (Panlab, Barcelona, Spain). The open field arena was divided into central (>8cm from any wall) and peripheral regions and homogeneously illuminated at 150 Lux. Each mouse was placed on the periphery of the open field and allowed to explore the apparatus freely for 30 min. The distance travelled, the number of rears and time spent in the central and peripheral parts of the arena were recorded over the test session by infrared beam breaks using a double infrared frame (multiple heights to capture the Z dimension) containing each a total of 16 x16 infrared cells at intervals of 2.5 cm.

#### Novel object recognition

The test was carried out in an open field arena as previously described(*9*). On the first and second days, mice were habituated to the arena for 15 minutes at 60 Lux. On the third day, animals were submitted to a 10-min acquisition trial during which they were individually placed in the presence of two of object A (a marble or die) placed 10 cm away from one of the box corners on the same side of the box. The exploration time of both object A (when the animal’s snout was directed towards the object at a distance ≤1 cm) was recorded by an observer from video. Then mice were then put back in their cage. After a retention period of 3 hours, a novel object discrimination test was conducted. One familiar object A and a novel object (object B, either marble or die, whichever was not used as object A) were placed at the same relative distance and position (approximately 20 cm) and the exploration time of these two objects over a period of 10 minutes was recorded by an observer from video. A discrimination index was defined as (tB/(tA + tB)) × 100.

##### Fear conditioning context discrimination

The procedure is adapted from (*30*) with polymodal operant chambers (Coulbourn Instruments, Allentown, PA) used for this experiment. We created two different environmental contexts: context A in chamber A was defined as a room lit with overhead lighting at 50 lux and containing two conditioning chambers. The 18.5 × 18 ×21.5 cm chamber had a plexiglass wall in front and back and aluminum side walls. The chamber floor consisted of a grid composed of 16 stainless steel rods connected via a cable harness to a shock generator. Context B in chamber B was different from context A in several ways: the overhead light location was changed, the wall was masked with black Plexiglas squares and the ceiling motif was changed. As in context A, the floor of each chamber consisted of 16 stainless steel rods which were wired to a shock generator and scrambler. The room was lit with a 30-W red overhead light.

On day 1 mice were placed in the context A conditioning room and into the conditioning chambers. After 4 minutes, they received a single unconditioned stimulus (US) (foot shock of 0.4mA for 1 second) and were removed from the chambers two min after foot shock termination. The next day, we subdivided each group of mice into two subgroups: 1 and 2. During the morning, subgroup 1 was tested in the same context as conditioning day (Context A) and subgroup 2 was tested in a new context (Context B). Each test consisted of 8min exposure to the chamber without the delivery of foot shock. The dependent measure employed was freezing behavior; the general activity of the animals was recorded through the infrared cell placed at the ceiling of the chambers directly on a PC computer using Graphic State (Coulbourn Instrument, Harvard Apparatus, Les Ulis, France). The behavior of each mouse was scored as freezing or not freezing every 2 seconds (with movement detected by the infrared sensors). These scores were then converted into a percentage of freezing. Freezing was recorded for four minutes in this context. During the afternoon (4-5 hours after the morning session), the contexts were reversed for each subgroup with subgroup 1 tested in the new context (context B) and subgroup 2 tested in context A.

#### Extraction of serum and brain for analysis of arbaclofen levels

Mice were euthanized with carbon dioxide and tissue was harvested on the day after the last behavioral test (day 31, after 29 days of arbaclofen treatment). Serum harvesting was performed 1 hour after lights-on and brain harvesting was performed 2 hours after lights-on, on the same day. Following isoflurane anesthesia, retro-orbitral blood (200 microliters) was collected in a Microvette 500 K3E (Sarstedt, Numbrecht, Germany). Serum was transferred to CryoTube Vials (1.8ML, Thermo Fisher, Illkirch, France) for storage at −80 °C. Brains were extracted, rinsed with saline, and bisected sagittally. Half of each brain was placed in a 15 mL conical vial and flash frozen to −80 °C.

### Abel Lab (University of Pennsylvania)

#### Ethical statement

All procedures were approved by the IACUC at the University of Pennsylvania.

#### Mice and housing conditions

Female B6129SF1/J (Jax lab stock# 101043) were crossed to B6129S-Del(7Slx1b-Sept1)4Aam/J mice (Jax lab stock# 013128)(*10*). Only male mice were used for this study. Upon weaning (P21-23), pups were housed in same-sex, mixed-genotype groups of 2-4 per cage and maintained in a temperature and humidity-controlled vivarium, with a 12 hr light/dark cycle (lights on at 07:00hr) and ad libitum access to food and drinking water. Mice were weighed weekly.

#### Drug treatment

Arbaclofen (R-baclofen) was provided as a generous gift from Clinical Research Associates. Arbaclofen was delivered *ad lib* in home cage drinking water at a concentration of 0.5mg/mL beginning at 7-9 weeks of age, for 12 days prior to, and throughout the duration of the behavior procedure battery (total 29 days). Consumption was measured and water bottles were changed daily.

#### Behavioral testing

The behavior battery consisted of an open field procedure (day 12 of arbaclofen treatment), novel object recognition (NOR) (days 14-24) and high speed rotarod (days 25-28). Behavioral testing was conducted in mice between 9 and 13 weeks of age. All behavior procedures were performed during the first four hours of the light phase. The order of testing the mice was randomized by cage (order of cages randomized, order of mice within each cage randomized). The investigator performing behavioral tests was blind to the genotype and treatment of each mouse. Mice were excluded from behavioral testing and analysis if they were deceased (1 wt,vehicle, 1 16p11.2 del,vehicle, 3 16p11.2 del,0.5 mg/kg arbaclofen). An additional cage of mice (4 mice, 2 wt, 2 16p11.2 del) were excluded from analysis because they were thought to be vehicle-treated but had appreciable levels of arbaclofen in brain tissue based on LC-MS/MS.

##### Open Field

Mice were allowed to habituate to the procedure room in their home cage for thirty minutes prior to the trial. Mice were allowed to explore the open field freely for 30 min. Spontaneous activity was measure in a 16” x 16” Plexiglas arena (ambient lighting 350 lux) fitted with a scaffold lattice of infrared emitters and detectors 1” apart (Photo activity System SDI, San Diego CA). Activity is detected as the mouse disrupts IR light beams. Horizontal activity (XY-axis), vertical activity (Z-axis), center activity (beam breaks >1 inch from the arena wall) and total activity (XY and Z-axis) were collected.

##### Novel Object Recognition

Mice were gently handled by the investigator performing the procedure for one to two minutes per day for five days before exposure to the arena. Mice were placed in an empty arena (lit to 275 lux) for five minutes per day for five consecutive days to allow for habituation to the arena. After five days of pre-exposure to the arena, mice were trained with a pair of the same objects for fifteen minutes, either two white sand-filled bottles (Michael’s Craft Shop) or two metal bars mounted vertically on a 2” x 2” acrylic base Twenty four hours later, a fifteen-minute recall trial was performed with one familiarized object explored during the training phase and a novel object (either bottle or bars, whichever was not used during training phase). Distance traveled during habitation and exploration time of objects was obtained during the training and recall phase using open source autophenotyping software(*31*). Exploration of the objects is defined as the time the snout of the mouse was within 2 cm of, and directed toward, the objects.

##### High Speed Accelerating Rotarod

Mice were allowed to habituate to the procedure room in their home cage for thirty minutes prior to the first trial. The Rotarod (IITC San Diego Ca.) was modified to attain speeds up to 80 rpm. Mice received three trials per session over four consecutive days as described in(*21*). Briefly, the first session starts with a two-minute habituation to the stationary rotarod before the initial low speed trial (accelerating 4-40 rpm/5 minutes). Two additional trials (4-40 rpm/5 minutes) were performed on day one. The next day, the mice received three low-speed trials. Three high speed trials (8-80rpm/5 minutes) were performed on days 3 and 4. All inter-trial intervals were about 30 minutes. Latency to fail was defined as the time to drop from the rod or time to make a full rotation while gripping on to the rod.

#### Extraction of serum and brain for analysis of arbaclofen levels

Mice were euthanized by cervical dislocation and tissue was harvested the day after the last behavior test (day 29) between 08:00-11:00hr to coincide with time of day of previous behavior testing. Trunk blood was collected in a tube with EDTA and placed on ice until centrifugation at 4 degrees C to separate plasma. Brain was removed, sagittally bisected, rinsed with 1-2 ml PBS, and then flash frozen in liquid nitrogen.

### Crawley Lab (University of California at Davis)

#### Ethical statement

All procedures were approved by the University of California Davis Institutional Animal Care and Use Committee.

#### Mice and housing conditions

16p11.2 deletion (*Coro1a-Spn*) mice, originally generated on a mixed C57BL/6N, 129P2/Ola, and CD-1 background by Thomas Portmann and Ricardo Dolmetsch at Stanford University(*11*) were re-derived at the University of California Davis and maintained on a mixed C57BL/6N, 129P2/Ola, and CD-1 background strain. Wildtype female mice were mated with heterozygous males to produce wildtype and heterozygous littermates. Both sexes were used for behavioral testing. Wildtype pups were weaned at 21 days of age and heterozygous mice were weaned at up to 35 days to reduce nutrition-related fatalities. Mice were housed in same-sex, mixed genotype groups of 2-4 littermates/cage, maintained in a temperature- and humidity-controlled vivarium, with *ad libitum* access to food and water on a 12 hr light/dark cycle (lights on at 07:00hr). Body weights were measured and recorded daily.

#### Drug treatment

Arbaclofen (R-baclofen) (provided as a gift from Clinical Research Associates) was administered in the home cage drinking water at a concentration of 0.5 mg/mL in Falcon 50ml conical centrifuge tubes fitted with drinking spouts. Treatment began at 7-9 weeks of age, 12 days before the start of behavioral testing, and continued throughout assay days until euthanasia for a total of 29 days. Amount of drinking water/arbaclofen consumed per cage was measured daily, using the grid lines on the Falcon tubes, and water/solution was changed daily.

#### Behavioral testing

The behavioral battery consisted of open field (day 12 of arbaclofen treatment), novel object recognition (days 14-16), and object location memory (days 19-21). Behavioral testing was conducted in male and female mice between 9 and 13 weeks of age. Behavioral tests were conducted during the light phase of the circadian cycle, at least one hour after lights on and one hour before lights off. On testing days, animals were placed in the experimental room antechambers 1 hour before the start of the experiment. The order of testing the mice was randomized by cage (order of cages randomized, order of mice within each cage randomized). The investigator performing behavioral tests was blind to the genotype and treatment of each mouse. Mice were excluded from behavioral testing and analysis in cases where an individual did not survive to the end of the testing sequence (1 wt,0.5 mg/ml arbaclofen, 2 16p11.2 del,0.5 mg/ml arbaclofen).

##### Open field

Mice were exposed to an empty AccuScan arena (40 × 40 × 30 cm, illuminated to 30 lux) for 30 minutes. Spontaneous activity (horizontal activity, rearing, and time spent in the center 20× 20 cm of the arena) was monitored by infrared beam breaks (8 × 8 × 2 infrared beams) using VersaMax Animal Activity software.

##### Novel Object Recognition

Novel object recognition testing was conducted in VersaMax Animal Activity Monitoring chambers (AccuScan Instruments). On days 1 and 2, each mouse was placed in an empty chamber (30 lux) and allowed to explore for 30 minutes, to habituate to the chamber. On day 3, the subject mouse was placed in the same empty chamber for a third habituation session of 10-minute duration. The mouse was removed, and two identical objects (plastic coral toys) were placed in opposite corners of the chamber. The mouse was then placed back into the chamber and allowed to explore the chamber containing the two identical objects for 10 minutes. After this familiarization session, mice were removed from their respective chambers and placed into separate holding cages in a different room for a one-hour inter-trial interval (ITI). Chambers were wiped down with ethanol between trials.

One of the objects from the familiarization session was replaced into the chamber in its original location. Where the second object had been, a novel and distinct object (a configuration of Duplo blocks) was placed. After the 1-hour ITI, the subject mouse was returned to the testing chamber and allowed to explore both objects for 5 minutes. Time spent sniffing each object was scored from videos using Noldus Ethovision three body point module with sniffing defined as the nose facing the object and ::: 2 cm from the object. To confirm equal salience of the two objects, the coral and Duplo objects were simultaneously presented to a previous group of 16p11.2 del mice. No innate object preference was detected.

##### Object Location Memory

Object location memory testing was conducted in VersaMax Animal Activity Monitoring chambers illuminated to 30 lux. On days 1 and 2, each mouse was placed in an empty chamber and allowed to explore for 30 minutes to habituate to the chamber. On day 3, the subject mouse was placed in the same empty arena for a third habituation session for 10 minutes. The mouse was removed, and 2 identical objects (toy treasure chests) were placed in the center of the chamber, ∼13 cm apart. The novel location was counterbalanced (left or right) across mice to prevent side bias. The mouse was then placed back into the chamber containing the two identical objects and allowed to explore for 5 minutes. After this familiarization session, mice were removed from the arena and placed into separate holding cages in a different room for a 1-hour inter-trial interval (ITI). Chambers were cleaned with ethanol between trials.

One of the objects from the training session was replaced in its original location, and the other object was moved to a new location, ∼18 cm forward from its initial location. After the 1-hour ITI, the subject mouse was returned to the testing chamber and allowed to explore both objects for 5 minutes. Time spent sniffing each object was scored from videos using Noldus Ethovision three body point module with sniffing defined as the nose facing the object and ::: 2 cm from the object.

#### Extraction of serum and brain for analysis of arbaclofen levels

Mice were euthanized and tissue harvested on day 28 of arbaclofen treatment between 3 and 6 hr after lights-on. Mice were anesthetized with 5% isoflurane and promptly decapitated. Trunk blood (∼ 300 microliters) was collected into BD K2EDTA Microtainer tubes. Tubes were inverted 10 times and centrifuged at 12.4 rpm for 3 minutes. Plasma was pipetted off the top (∼ 100 microliters) into Nalgene Thermo 100 cyrogenic tubes and stored at −80°C. Brains were rapidly removed from skull, rinsed with saline, dissected along the sagittal midline, and flash frozen in a slurry of acetone and dry ice. Brain halves were stored at −80°C.

### Datta Lab (Harvard University)

#### Ethical Statement

All experiments were approved by the Institutional Animal Care and Use Committee at Harvard Medical School (protocol number IS00000138).

#### Mice and housing conditions

Heterozygous 16p11.2 deletion (*7Slx1b-Sept1*) mutants (16p11.2df, stock No: 013128)(*10*) and wild type mice (B6129SF1/J, stock No: 101043) were obtained from the Jackson Laboratory. Mice used in this study were offspring of heterozygous mutant and wild type breedings (both female het x male wt and male het x female wt). Only male mice were used in this study. Mice were weaned from P21-P28 and housed in same-sex, mixed-genotype groups of 2-5 mice/cage. Mice were transferred to the reverse light cycle room (12 hour light/dark, lights on at 22:00hr) at least one day before the start of drug administration to perform the behavior experiments in the dark cycle. Mice were weighed daily during the drug administration.

#### Drug administration

##### Arbaclofen

Arbaclofen (a gift of Clinical Research Associates) was dissolved in tap water to a concentration of 0, 0.25, 0.5 or 1.0 mg/ml and provided *ad lib* to each cage in regular water bottles. Drug administration began at 8 weeks of age, 12 days before the start of behavioral testing and continued for a total of 29 days. Bottles were changed daily and filled with freshly prepared solution.

We we focused our detailed analyses on 0, 0.5, 1.0 mg/mL doses to examine the effects of arbaclofen as we did not observe reproducible effects that were distinct from control in the 0.25 mg/mL condition.

##### Risperidone

Risperidone (Sigma, 106266-06-2) dissolved in saline (lactated ringer) at a dose of 0.2 mg/kg was administered to mice by daily IP injection from day 1 to day 13. The injection on day 13 was performed 1 hour before behavioral recording.

##### Behavioral testing

For the arbaclofen experiment, the mice were exposed to the open field on day 13 of arbaclofen administration, with drug administration continuing until day 29. For the risperidone experiment, mice were exposed to the open field on day 13 of drug treatment. Behavioral testing was conducted in mice at 10 weeks of age. All experiments were performed during the dark phase of the reverse day-night cycle. Mice were habituated to the procedure room for 10 minutes in their home cage before the start of behavioral testing. The order of testing the mice was randomized by cage (order of cages randomized, order of mice within each cage randomized). The investigator performing behavioral tests was blind to the genotype of each mouse, as well as to risperidone vs. vehicle treatment (but not arbaclofen vs. vehicle treatment). Mice were excluded from analysis if they were not within the range of 21.3g ∼ 29.5g, which spans ∼2× standard deviation of all mice. 4 animals died or euthanized during the experiment due to poor health (2 0.5 mg/ml arbaclofen and two 1.0 mg/ml arbaclofen, genotype unknown).

##### Open field

Open field testing began on day 13. Each mouse was manually placed into a square, matte acrylic black box (bottom: 40 cm x 40 cm, height: 30 cm) and recorded with a kinect2 camera (Microsoft) at 30 fps for 15 minutes, taken out of the box and returned to their home cage (still in the procedure room) for 45 minutes, and then imaged for a second 15-minute session.

#### Data pre-processing

Depth data were extracted by custom software. This yielded conventional scalar data describing mouse behavior, including the position of centroid, length, height, width, angle and velocity of the mouse. In addition, the image of the mouse was extracted and oriented from the open field movies, generating a new movie in which the mouse was centered in an 80×80 pixel box, with its nose digitally aligned towards the right, and its tail towards the left. The tail was then removed via filtering. This extracted, centered and oriented movie was used as the substrate for all further modeling.

#### Motion Sequencing analysis

Motion Sequencing (MoSeq) is a catch-all term for a combined machine vision and machine learning system that automatically identifies behavioral motifs and the order in which they occur(*20*). The MoSeq algorithm first takes the aligned and centered 80 x 80 pixel aligned movies of each mouse and performs PCA on this high-dimensional datastream to lower the dimensionality of the data that is being analyzed. Note that this PCA procedure is only performed to make the subsequent computations easier; MoSeq returns similar results when fed raw pixels, although at significant computational cost. Here the first 10 principal components were used, which capture approximately 90 percent of the overall variance in the data.

These PCA-reduced 3D imaging data are then fit to a generative model for mouse behavior using computational inference techniques. The model and fitting procedure use regularities in the data to automatically discover, in any behavioral dataset, the optimal number of behavioral syllables, the identity of each of these behavioral syllables (in terms of the specific pose trajectories that define each syllable) and the frequency with which each syllable transitions from one to the other. MoSeq also labels the dataset that was used for training: for each frame of 3D data, MoSeq identifies the most likely behavioral syllable to be expressed. Thus, MoSeq takes as its input 3D imaging data of mice, and returns a set of behavioral syllables that characterizes the expressed behavior of those mice, and the statistics that govern the order in which those syllables were expressed in the experiment. We used MoSeq2 with robust AR-HMM that use student’s t-distributed AR-HMM model, kappa = 2e5, nu = 10, iter = 500. Details of the MoSeq analysis pipeline were mentioned in (*20*) and (*32*).

#### Analysis of behavioral syllables

Syllables identified by MoSeq were numbered based upon usage frequency, which was calculated by the fraction of frames accounted for by each syllable, across all experimental groups and usages for each group. The 36 most-used syllables each appeared in >1% of total frames and collectively they accounted for 94.4 % of total frames. We excluded syllable 21 as it was an obvious noise syllable, and thus used 35 syllables for the downstream analyses. For analyzing the transition probability between syllables, we calculated the bigram probabilities as below. The Bigram probability between syllable i and j (Pij) is defined as the number of transitions from syllable i to j normalized by the number of total transitions. Cytoscape was used to visualize the bigram probability matrix. We only used the first trial of each session as the behavior of the second trial was less reliable presumably due to handling stress.

#### Extraction of serum and brain for analysis of arbaclofen levels

Mice were euthanized with carbon dioxide and tissue harvested on day 29 at 1 hr after lights on (23:00hr). Blood was collected and transferred into a BD microtainer (BD, 365974) and centrifuged for 3 min (5000xg); supernatant was transferred to cryotubes (Thermo, 5000-1020) and flash frozen. Brains were dissected, bisected sagitally, rinsed with lactated ringer, transferred to a cryotube, and frozen on dry ice. Both serum and brains were stored at −80C until shipment.

### All four labs in the consortium

#### Analysis of arbaclofen levels in mouse brain

Analysis of arbaclofen levels in mouse brain was performed by Pace Analytical Life Sciences (Woburn, MA).

Mouse brain homogenate was analyzed for arbaclofen content following a liquid-liquid extraction via LC-MS-MS. 30% (w/v) mouse brain homogenate was prepared by weighing pre-portioned mouse brain into 1.5 mL microcentrifuge tubes, adding 0.1% formic acid in water (v/v) at a volume 2.33 times the weight of the mouse brain portion, and homogenizing for 10-20 seconds with a hand-held homogenizer.

Calibration standards (0.5-1000 mg/mL) were prepared from a 0.5 mg/mL arbaclofen stock solution in 50/50 water/methanol. To produce a full matrix match between standards and test samples, calibration standards were diluted with blank 30% mouse brain homogenate acquired from Bio IVT at a ratio of 10 µL arbaclofen stock to 150 µL of mouse brain homogenate and for test samples 10 µL 50/50 water/methanol was added to 150 µL of each test sample. All calibration standards, appropriate QC samples and blanks, and test samples were then extracted with 600 µL of acetonitrile spiked with an internal standard (arbaclofen-d4) at 100 ng/mL. the extracted samples and standards were mixed by orbital shaking at 700 rpm for 5 mins and then centrifuged at 3220 rcf for 5 minutes. The supernatants were transferred to a 96-well plate and dried completely under nitrogen at 60C. The dried samples were reconstituted in 100 μL of water/methanol (50/50) for analysis by LC-MS.

The LC-MS method used to analyze the extracted samples consisted of a Shimadzu LC-20AD HPLCS, CTC HTS PAL Autosampler and an ABI/MDS Sciex API 4000 Mass Spectrometer running Analyst 1.6.2 software. A C18 column (Mac-Mod ACE 3 C18, 2.1 x 50 mm, 3 um P/N: ACE-111-0502) was used. Mobiles phases were 2 mM ammonium acetate with 0.1% formic acid in 95:5 and 5:95 water:methanol for MPA and MPB respectively. The mobile phase gradient ran for four minutes per injection at 0.5 mL/min: 10% MPB to 100% MPB from 0.5-3 minutes followed by 1 minute at starting conditions (10% MPB). Column temperature was kept at ambient and sample temperature was 10°C. The injection volume was 20 µL. The transitions monitored were 214.2→151.0 for arbaclofen and 218.2→155.1 for arbaclofen-d4.

Arbaclofen content in the mouse brain samples was calculated using a curve generated by the calibration standards and corrected for the dilution factor introduced with the homogenization of the brain samples.

#### Statistical analysis

All datasets met the assumptions of parametric tests of normally distributed data and equal variances across groups. Outliers (data points > +/-2 standard deviations from the mean) were excluded from analysis. For all tests, alpha was set to 0.05.

##### Novel Object Recognition/Object Location Memory

Mice with total exploration times (for both objects) less than 3 seconds in either training or testing phase were excluded from analysis. Raw exploration time data were used to calculate a discrimination index of percent time exploring novel object (or object in novel location) divided by the total time spent exploring both objects during the recall phase. Discrimination index data were analyzed by 2-way ANOVA (genotype and treatment as between-subjects factors) with post-hoc Welch’s t-tests for individual group comparisons (adjusted for multiple comparisons by Holm-Sidak method). Tests were conducted in GraphPad Prism v8 for MacOS. Raw exploration time data were analyzed by 3-way RMANOVA (object or location as within-subjects variable, genotype and treatment as between-subjects variables), conducted with ‘aov’ in r.

##### Contextual fear conditioning

Raw freezing scores were converted to percentage freezing (time spent freezing divided by total time). Percentage freezing data were analyzed by 3-way RMANOVA (context as within-subjects variable, genotype and treatment as between-subjects variables), conducted in Sigma Plot 11.0.

##### Rotarod

Latency to fall data were analyzed by 3-way RMANOVA (time as within-subjects variable, genotype and treatment as between-subjects variables), conducted in SPSS (IBM version 24).

##### Open field

Percent time spent in center was calculated as % time spent in center = 100% x (time spent in center/total time). Activity count data were analyzed by 3-way RMANOVA (time as within-subjects variable, genotype and treatment as between-subjects variables), conducted with ‘aov’ in r.

##### MoSeq open field analysis

To analyze syllable usage data, we performed z-test with variance estimated by 1000 bootstraps and p-values were corrected by Benjamini-Hochberg FDR as in (*20*). We reported the significant syllables if corrected p-value was < 0.05.

For the bigrams/trigrams, bigrams/trigrams detected more than chance were identified by Monte Carlo randomization for each condition (50000 times:p < 5e-5 for bigrams. 100000 times:p<2e-5 for trigrams, one sided). The union of such bigrams/trigrams in different groups was then compared by bootstrap test (1000 times). Significantly different bigrams/trigrams were identified if 95% confidence intervals were not overlapped. P-values for the summed difference were calculated as follows:

p = (# of bootstraps where the value for group A is less than that for group B + 1) / (# of bootstraps)

p values were subjected to Bonferonni correction, and corrected p values <0.05 were considered significant.

## Acknowledgements

General

We thank Sarah Fletcher, Carina Luck and Philipp Schneggenburger at Pace Analytical Life Sciences for performing the analysis of arbaclofen content in mouse brain tissue. We thank Bing Han at the Simons Foundation for help with statistical analyses. We thank Emalee Peterson, Ralph E Peterson, and Paneed Jalili for help with the experiments (both behavior and tissue sampling) in the Datta lab.

## Author contributions

BBG, ALC, TA, JNC, SRD, and YH conceptualized the study. BBG performed formal analysis of data for this study. The original draft of this manuscript was generated by BBG. Methodology was prepared and investigation was carried out by SML and YH for the IGBMC partner with internal validation of data. In the Crawley lab, MDS and MS performed the behavioral testing, harvested the tissues, and contributed to writing the manuscript; JNC designed and supervised the experiments and contributed to writing the manuscript. In the Abel lab, WTO designed and performed the behavior experiments and tissue harvest. In the Datta lab, data generation and analysis were performed by TT and SRD, and funding acquisition and supervision was performed by SRD. The paper was reviewed and edited by all authors.

## Data and material availability

Data are available upon request to the authors.

## Supplemental Figures

**Figure S1.**
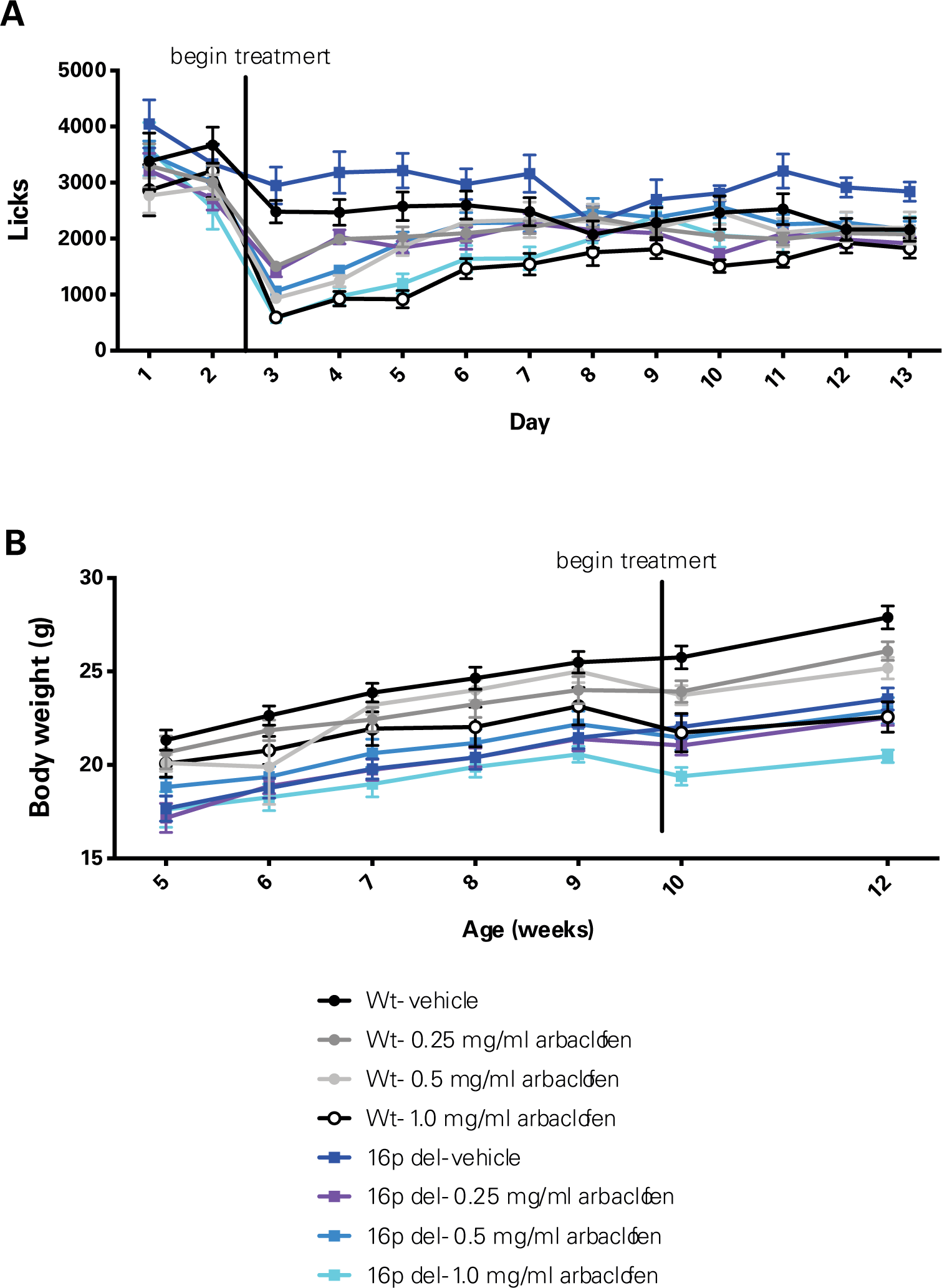
Arbaclofen administered in drinking water temporarily reduces drinking and body weight. **A-B.** Data were collected in the Herault lab (Del6 model). **A.** Mice were housed in intellicages for 2 days prior to the start of and 12 days of arbaclofen treatment. The number of licks at water bottles for mice of each genotype is shown. Data were analyzed by 3-way RMANOVA (genotype and treatment as between-subjects factors, day as within-subjects factor). 3-way RMANOVA: significant main effect of treatment (F(3, 95) = 12.674, p = 4.88 x 10^-7^), and day (F(12, 95) = 53.323, p< 2×10^-16^), no main effect of genotype (F (1, 95) = 2.821, p = 0.0963). Significant genotype x day interaction (F(12, 95) = 1.761, p = 0.0499) and treatment x day interaction (F(36, 95) = 5.494, p<2×10^-16^. Mean +/-SEM is graphed. n = 14, 14, 13, 10, 14, 13, 16, 9. **B.** Mice were weighed weekly. Body weight (g) is shown. Data were analyzed by 3-way RMANOVA (genotype and treatment as between-subjects factors, week as within-subjects factor). 3-way RMANOVA: significant main effects of genotype (F(1, 70)= 48.829, p = 1.31×10^-9^), treatment (F(3, 70) = 3.358, p = 0.0236), and week (F(6, 70) = 141.012, p< 2×10^-16^). Significant treatment x week interaction (F(18, 70) = 4.184, p=3.63×10^-8^). Mean +/-SEM is graphed. n = 10, 10, 10, 7, 11, 11, 11, 9.

**Figure S2.**
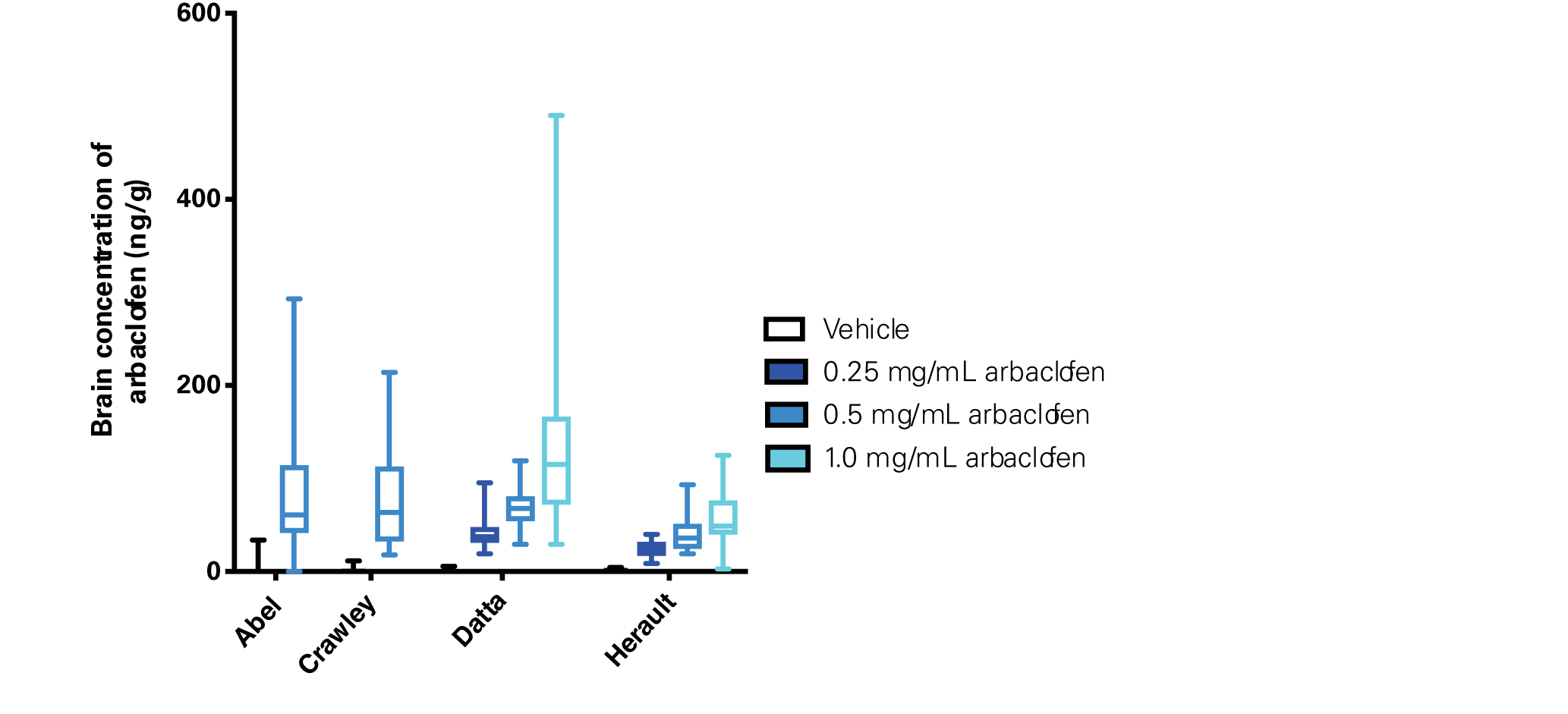
Arbaclofen is present in brain tissue of treated wildtype and 16p11.2 deletion model mice in dose-dependent concentrations. Brains were harvested from wildtype and 16p11.2 deletion model mice at the conclusion of behavioral studies, following 29 days of treatment. HPLC-MS/MS was used to analyze arbaclofen concentrations in frozen brain tissue, plotted here by study site. Graphs are box and whisker plots; whiskers are maximum and minimum, boxes are interquartile range, and midline is median. n = 21, 26, 34, 33, 38, 36, 41, 37, 30, 28, 29, 25

**Figure S3.**
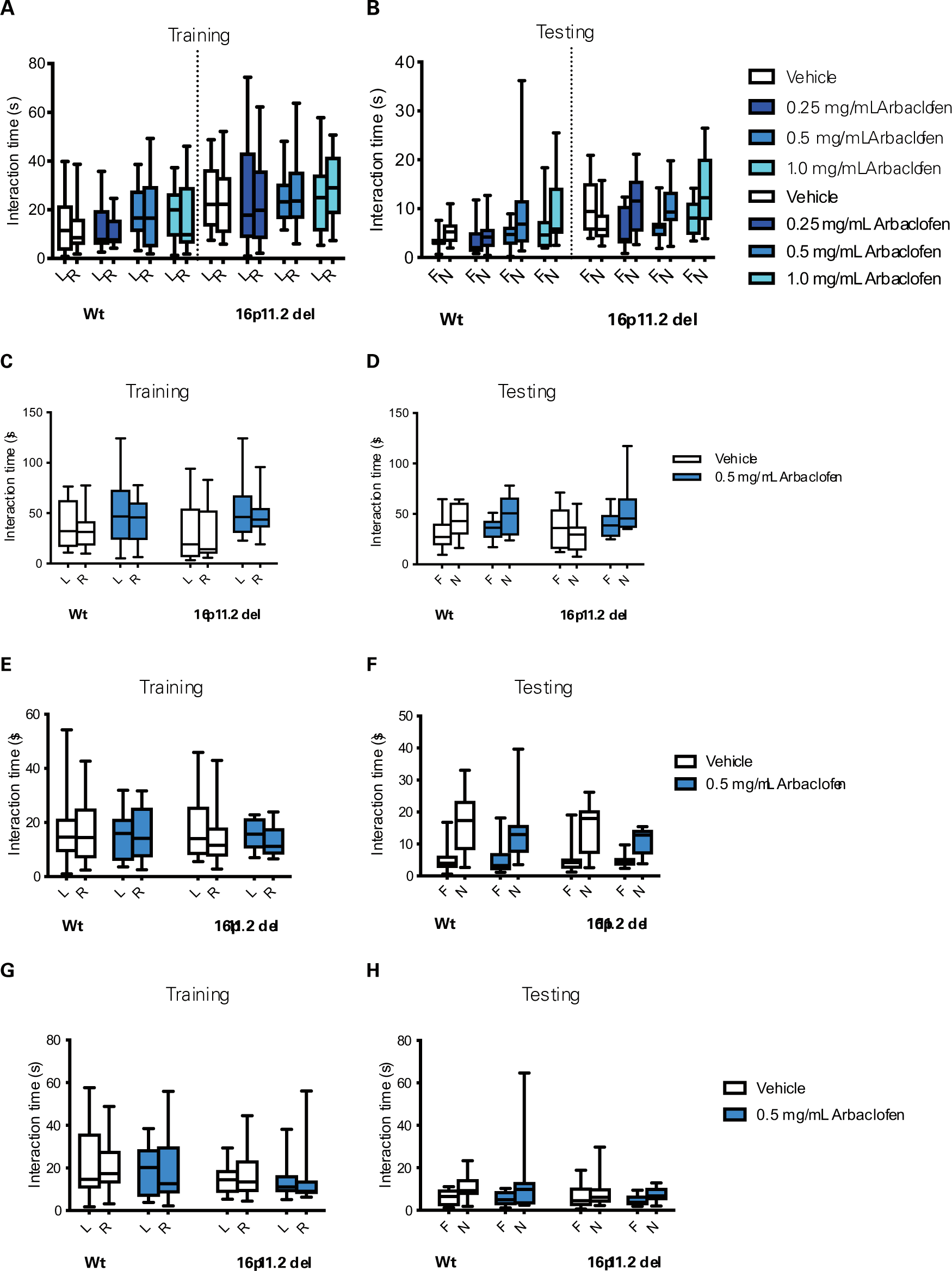
Arbaclofen rescues recognition memory deficits observed in two models of 16p11.2 deletion mice. Graphs are box and whisker plots; whiskers are maximum and minimum, boxes are interquartile range, and midline is median. L= left object; R= right object; F= familiar object or location; N= novel object or location. **A-F.** During the training phase of the novel object recognition task, wt and 16p11.2 del model mice treated with vehicle or one of 3 doses of arbaclofen were allowed to interact with two identical objects. After a delay, mice were tested by exposure to one of the familiar objects and one novel object. Time spent exploring each object during training (A, C, E) and testing (B, D, F) phases is shown. Data were analyzed by 3-way ANOVA (genotype and treatment as within-subjects factors, object as within-subjects factor. **A,B:** Data collected in the Herault lab (Del6 model; 3-hour delay between training and testing). n= 14, 13, 13, 11, 13, 13, 14, 12. **A.** Training phase. 3-way RMANOVA: significant main effect of genotype (F(1, 95) = 17.474, p = 6.48x 10^-5^), no significant main effect of object (F(1, 95) = 0.182, p = 0.670), treatment (F(3, 95) = 0.805, p = 0.494), or interactions. **B.** Testing phase: 3-way RMANOVA: significant main effect of genotype (F(1, 95) = 17.234, p=7.21×10^-5^), significant main effect of object (F(1, 95) = 25.974, p=1.76×10^-6^), no significant main effect of treatment (F((3, 95) = 2.496, p=0.0645). Significant genotype x treatment x object interaction (F(3, 95) = 3.488, p = 0.01875). **C,D:** Data collected in the Abel lab (Del4 model; 24-hour delay between training and testing). n= 16, 14, 7, 12. **C.** Training phase: 3-way RMANOVA: No significant main effects of genotype (F(1, 45) = 0.211, p=0.6478), treatment (F(1, 45) = 3.771, p = 0.0584), or object (F(1, 45) = 3.799, p = 0.0575). No significant interactions. **D.** Testing phase: 3-way RMANOVA: significant main effect of object (F(1, 45) = 16.284, p = 0.000209), no significant main effect of treatment (F(1, 45) = 3.939, p=0.0533) or genotype (F(1, 45) = 0.186, p = 0.6681). Trend toward genotype x treatment x object interaction (F(1, 45) = 2.984, p = 0.0909). **E. E,F:** Data collected in the Crawley lab (Del1 model; 1-hr delay between training and testing). n= 18, 17, 14, 10 **F.** Training phase: 3-way RMANOVA: No significant main effects of genotype (F(1, 55) = 0.45, p =0.832), treatment (F(1, 55) = 0.379, p = 0.541), or object (F(1, 55) = 0.704, p = 0.405). No significant interactions. **G.** Testing phase: 3-way RMANOVA: Significant main effect of object (F(1, 55) = 89.456, p = 3.95×10-^13^). No significant main effects of genotype (F(1, 55) = 0.831, p = 0.371), treatment (F(1,55)=1.911, p=0.172), or significant interactions. **G-H.** During the training phase of the object location memory task, wt and 16p11.2 Del1 model mice treated with vehicle or a single dose of arbaclofen were allowed to interact with two identical objects. After a delay, mice were tested by exposure to the two trained objects, one in its previous location, and one moved to a novel location within the testing arena. Time spent exploring each object during training (G) and testing (H) phases is shown. Data collected in the Crawley lab. Data were analyzed by 3-way ANOVA (genotype and treatment as within-subjects factors, location as within-subjects factor). n= 15, 17, 14, 12. **G.** Training phase: 2-way RMANOVA: No significant main effects of object (F(1, 54) = 0.144, p =0.706), genotype (F(1,54) = 2.969, p = 0.0906), treatment (F(1,54) = 0.267, p =0.6072), or significant interactions. **H.** Testing phase: 3-way RMANOVA: significant main effect of location (F(1, 54) = 12.376, p = 8.9×10^-4^). No significant main effects of genotype (F(1,54) = 0.949, p = 0.334), treatment (F(1,54) = 0.099, p = 0.754), or significant interactions.

**Figure S4.**
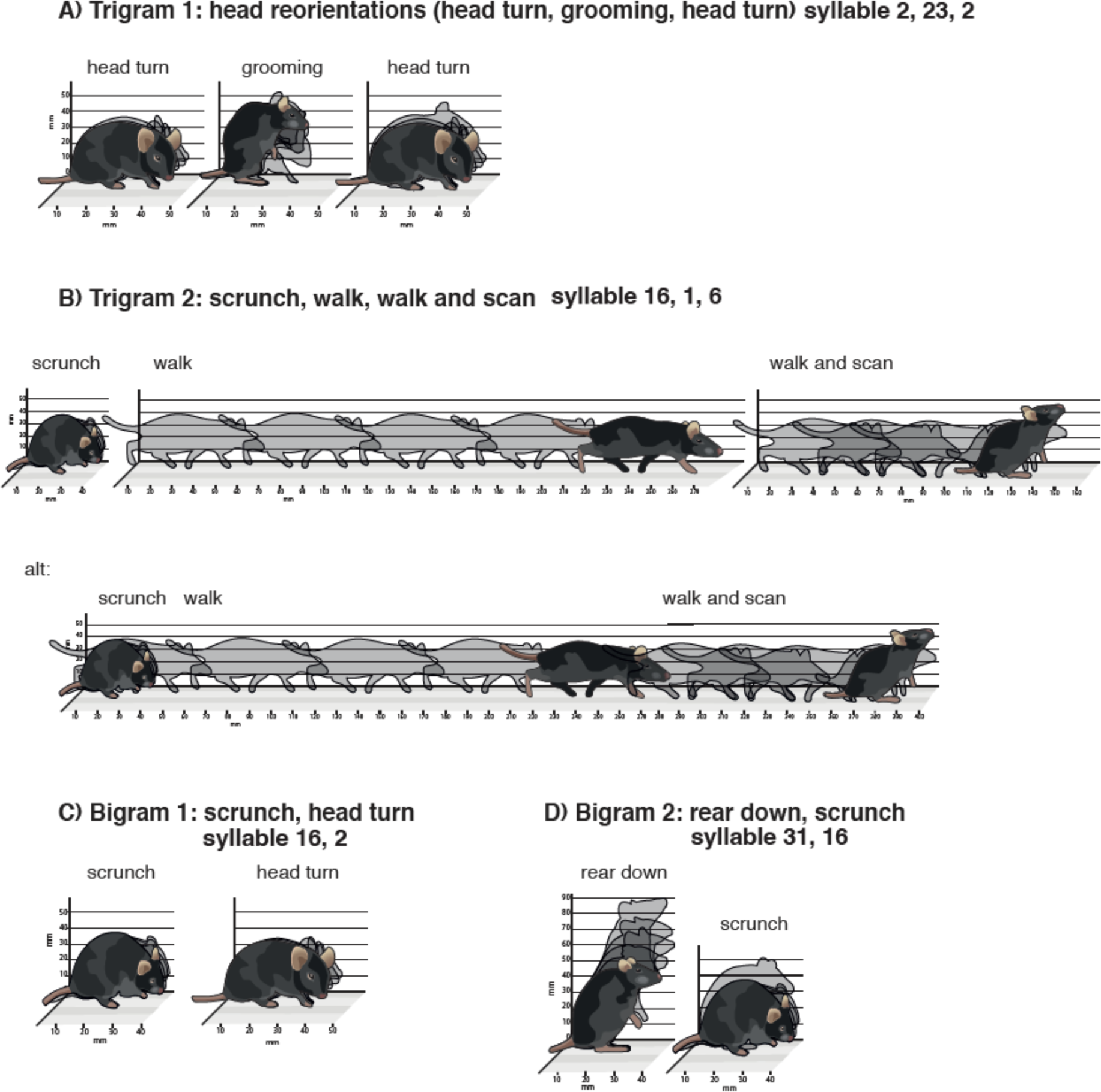
Artist’s rendering of bigrams and trigrams. Figure credit: Sigrid Knemeyer, Scistories.

**Figure S5.**
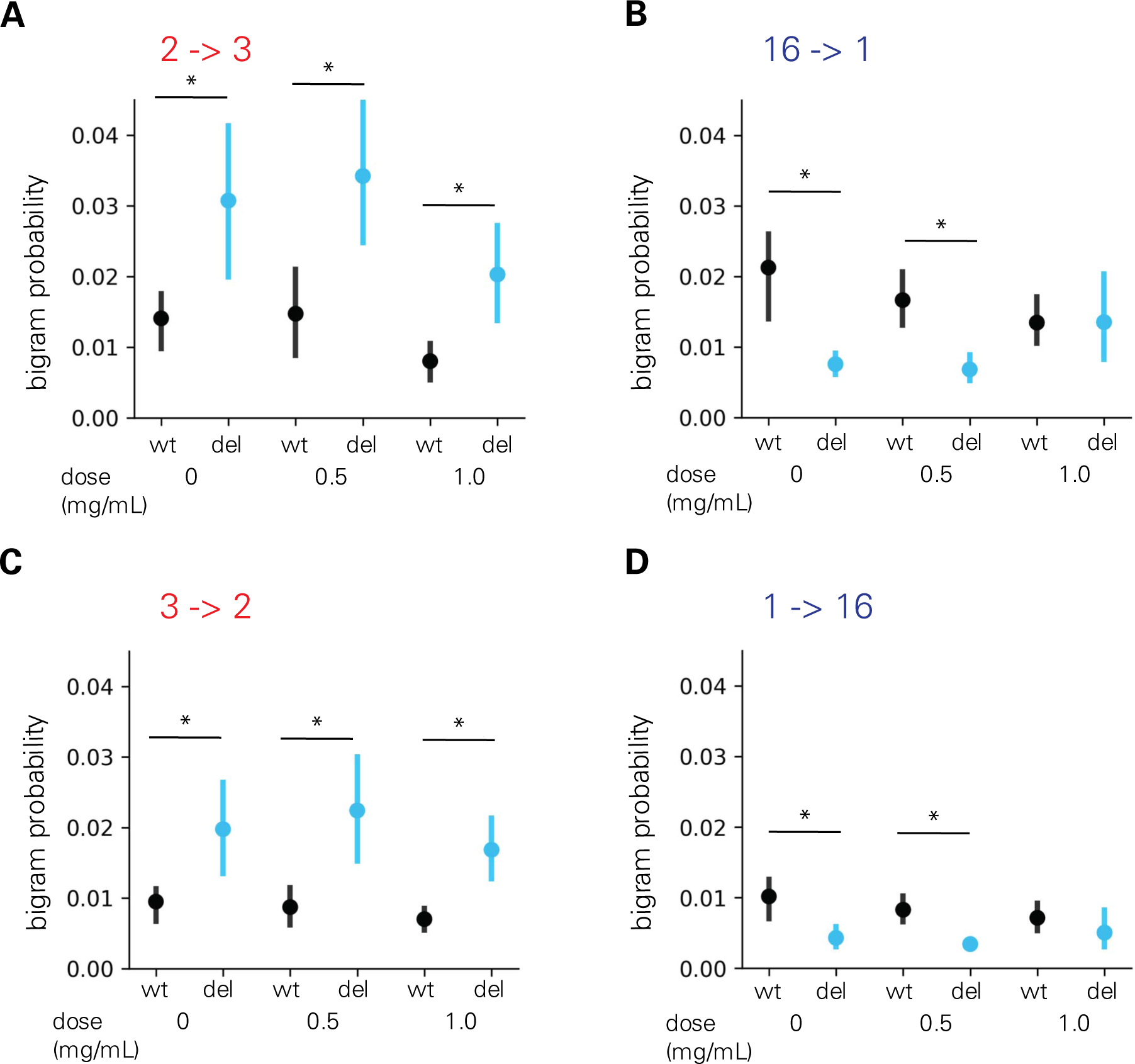
Differences in bigram probability in 16p11.2 Del4 deletion model mice are rescued by arbaclofen treatment. **A-D.** Data collected in the Datta lab. Example graphs showing probabilities of expressing individual bigrams across genotype and treatment groups. Error bars indicate 95% confidence intervals from 1000 bootstrapping and asterisks indicate non-overlap of confidence intervals between wildtype and 16p11.2 del for each condition. n= 15 wt/vehicle, 12 16p del/vehicle, 27 wt/0.5 mg/mL arbac, 14 16p del/0.5 mg/mL arbac, 18 wt/1.0 mg/mL arbac, 11 16p del/1.0 mg/mL arbac

**Figure S6.**
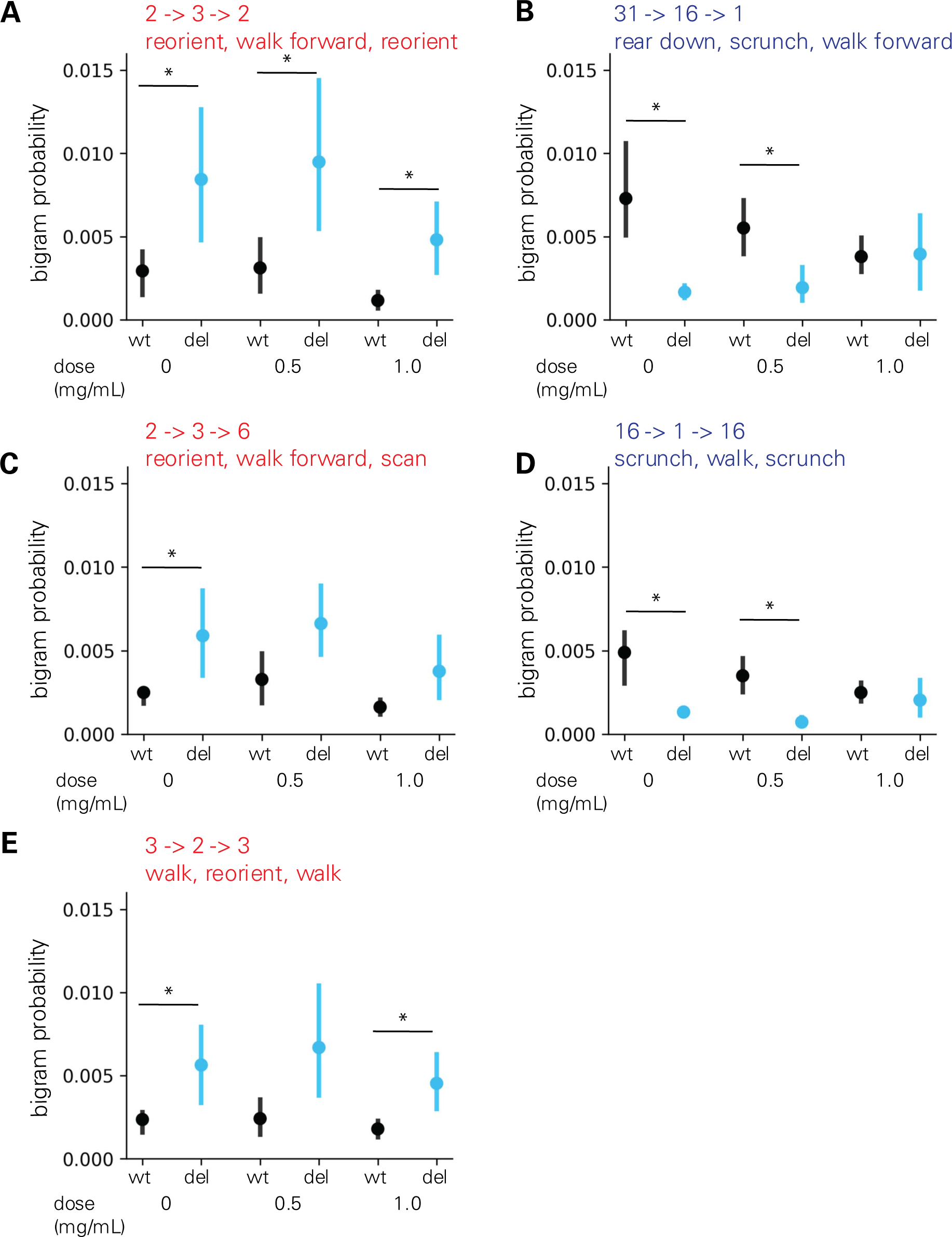
Differences in trigram probability in 16p11.2 Del4 deletion model mice are rescued by arbaclofen treatment. **A-E.** Data collected in the Datta lab. Example graphs showing probabilities of expressing individual trigrams across genotype and treatment groups. Error bars indicate 95% confidence intervals from 1000 bootstrapping and asterisks indicate non-overlap of confidence intervals between wildtype and 16p11.2 deletion model mice for each condition. n= 15 wt/vehicle, 12 16p del/vehicle, 27 wt/0.5 mg/mL arbac, 14 16p del/0.5 mg/mL arbac, 18 wt/1.0 mg/mL arbac, 11 16p del/1.0 mg/mL arbac

**Figure S7.**
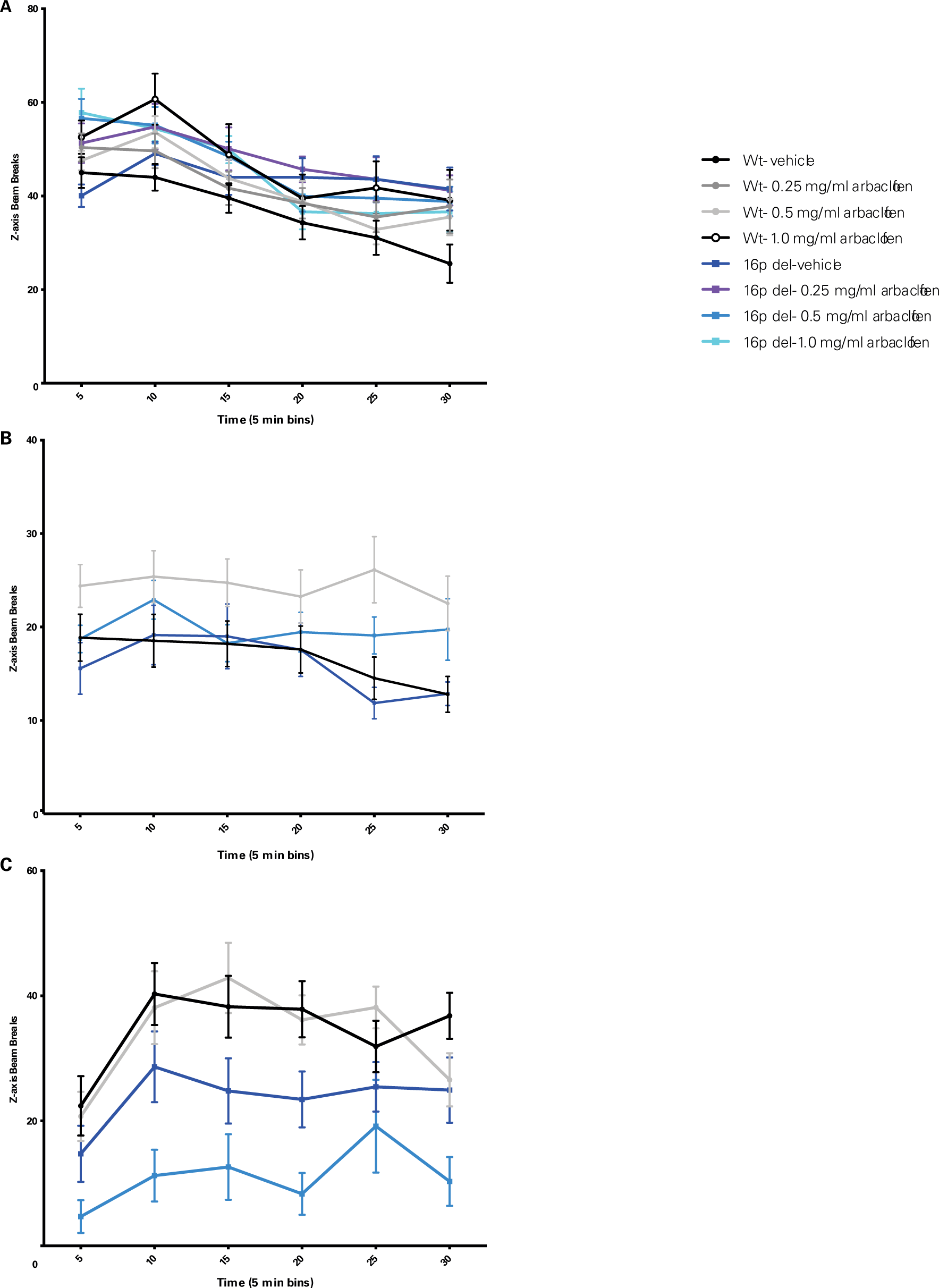
Inconsistent effects of arbaclofen on rearing behavior. Wildtype and 16p11.2 deletion model mice treated with one of 3 doses of arbaclofen were exposed to an open field for 30 min. Rearing counts (measured by infrared beam breaks in the z plane) are plotted in 5-minute bins. Points are means with error bars indicating SEM. Data were analyzed with 3-way RMANOVA (genotype and treatment as between-subjects variables, time as within-subject variable). **A.** Data were collected in the Herault lab (Del6 model). 3-way RMANOVA: main effect of time (F(5,101) = 50.699, p<0.0001), no significant main effect of genotype (F(1, 101) = 2.026, p = 0.1577), or treatment (F(3, 101) = 1.154, p = 0.3312). Significant time x treatment x genotype interaction (F(15, 101) = 2.067, p = 0.0103). n = 14, 14, 14, 12, 14, 14, 15, 11. **B.** Data were collected in the Abel lab (Del4 model). 3-way RMANOVA: significant main effect of time (F(5, 44) = 3.092, p=0.0102), significant main effect of treatment (F(1, 44) = 7.824, p=0.00762), no significant main effect of genotype (F(1, 44) = 1.044, p = 0.31245). Significant time x treatment interaction (F(5, 44) = 2.265, p=0.0491). n = 15, 15, 7, 11. **C.** Data were collected in the Crawley lab (Del1 model). 3-way RMANOVA: significant main effect of time (F(5, 64) = 14.080, p = 1.87×10^-12^), significant main effect of genotype (F(1, 64) = 17.712, p = 8.18 × 10^-5^), no significant main effect of treatment (F(1, 64) = 2.037, p = 0.158). Significant time x treatment interaction (F(5, 64) = 2.420, p=0.0357). n = 20, 21, 14, 13.

**Figure S8.**
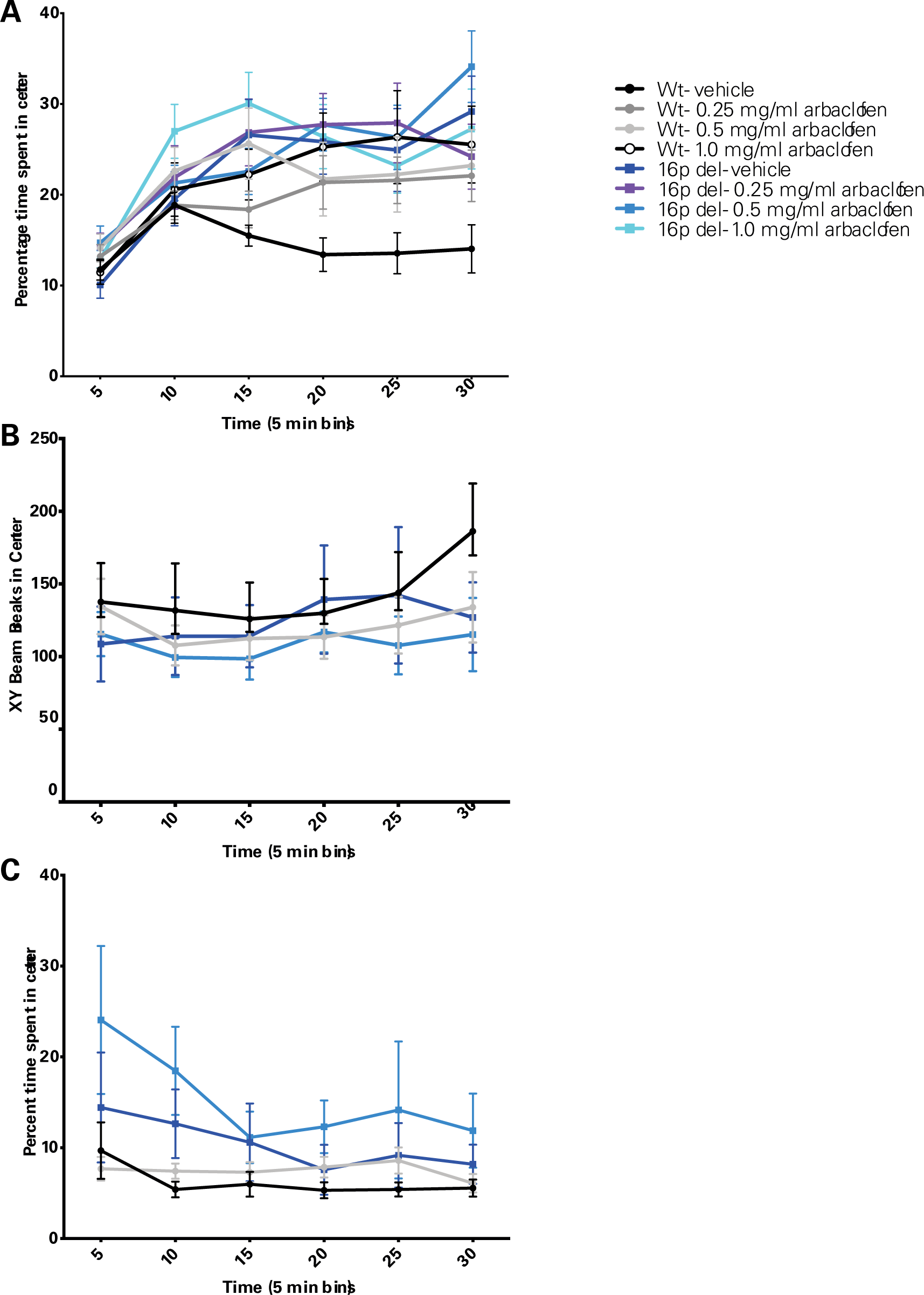
Inconsistent effects of arbaclofen on behavior in the center of the open field. Wildtype and 16p11.2 deletion model mice treated with one of 3 doses of arbaclofen were exposed to an open field for 30 min. Distance traveled in the center of the open field (measured as infrared beam breaks in the x and y planes within a predefined square in the center of the larger arena) or percentage of time spent in the center of the open field is plotted in 5-minute bins. Points are means with error bars indicating SEM. Data were analyzed with 3-way RMANOVA (genotype and treatment as between-subjects variables, time as within-subject variable). **A.** Data were collected in the Herault lab (Del6 model). Percent time spent in the center of the open field is plotted. 3-way RMANOVA: significant main effect of time (F(5, 101) = 33.454, p < 0.0001), significant main effect of genotype (F(1,101) = 6.979, p = 0.0096), no significant main effect of treatment (F(3,101) = 1.650, p = 0.1827), significant time x genotype interaction (F(5,101)=3.211, p =0.0073), significant time x genotype x treatment interaction (F(15, 101) = 2.404, p = 0.0022). n = 14, 14, 14, 12, 14, 14, 15, 12. **B.** Data were collected in the Abel lab (Del4 model). Distance traveled in the center of the open field is plotted. 3-way RMANOVA: significant main effect of time (F(5, 45) = 4.082, p=0.00144). No significant main effect of treatment (F(1, 45) = 1.087, p = 0.303), genotype (F(1, 45) = 0.802, p = 0.375), or significant interactions. n = 16, 14, 8, 11. **C.** Data were collected in the Crawley lab (Del1 model). Percent time spent in the center of the open field is plotted. 3-way RMANOVA: significant main effect of time (F(5,61) = 2.559, p=0.0275), significant main effect of genotype (F(1,61) = 9.169, p = 0.0036), no significant main effect of treatment (F(1,61) = 1.697, p = 0.1976). No significant interactions. n = 21, 20, 14, 12.

**Figure S9.**
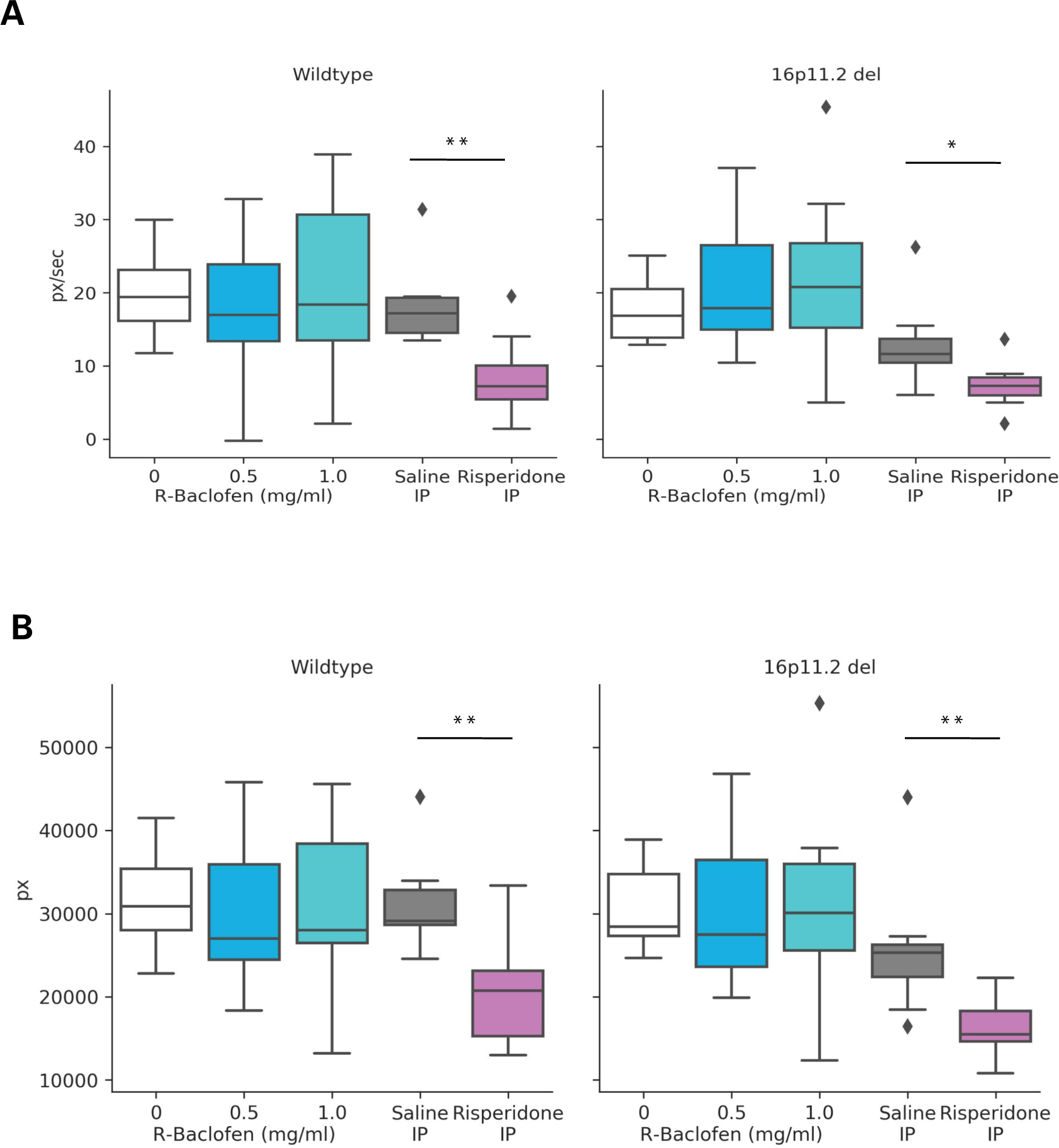
Risperidone reduces velocity of movement and distance traveled regardless of genotype, whereas arbaclofen does not affect these measures. Data were collected in the Datta lab (Del4 model). Graphs are box and whisker plots; whiskers are maximum and minimum, boxes are interquartile range, and midline is median. Wildtype: n= 15, 27, 18, 6, 13. 16p11.2 del: n= 12, 14, 11, 11, 8. *p < 0.05 vs. vehicle; **p < 0.01 vs. vehicle. **A.** Mean angular velocity over the entire recording session was measured from videos. Mann-Whitney U-test was performed to examine the difference between control and drug groups (separate analysis for arbaclofen and risperidone vs. control). Wildtype: 0.5 mg/mL arbaclofen vs. vehicle U = 171, p = 0.4158. 1.0 mg/mL arbaclofen vs. vehicle: U = 131, p = 0.8993. Risperidone vs. vehicle: U = 6, p =0.0044. 16p11.2 del: 0.5 mg/mL arbaclofen vs. vehicle U = 74, p = 0.6251. 1.0 mg/mL arbaclofen vs. vehicle: U = 47, p = 0.2549. Risperidone vs. vehicle: U = 15, p =0.0186. **B.** Total distance traveled over the entire recording session was measured from videos. Mann-Whitney U-test was performed to examine the difference between control and drug groups (separate analysis for arbaclofen and risperidone vs. control). Wildtype: 0.5 mg/mL arbaclofen vs. vehicle U = 166, p = 0.3447. 1.0 mg/mL arbaclofen vs. vehicle: U = 117, p = 0.8993. Risperidone vs. vehicle: U = 8, p =0.0075. 16p11.2 del: 0.5 mg/mL arbaclofen vs. vehicle U = 70, p = 0.4875. 1.0 mg/mL arbaclofen vs. vehicle: U = 66, p = 0.9755. Risperidone vs. vehicle: U = 6, p =0.0020.

**Movie S1. syllable16-2_bigram_scrunch-turn.mp4**

This “crowd” movie is generated by collating and overlaying 20 individual movies of a bigram (sequence of syllable 16 and 2) from multiple animals in wt and 0 mg/mL condition into a single movie. Green dots on mice indicate they are executing the bigram. Colors represent the height of any given pixel above the floor of the arena and the “cubehelix” from python’s matplotlib color maps was used (black, green, pink, white, from low to high). Note that these crowd movies are consistent across genotypes and doses.

**Movie S2. syllable31-16_bigram_reardown-scrunch.mp4**

Same as Supplementary movie 1, but for another bigram (sequence of syllable 31 and 16).

**Movie S3. syllable2-23-2_trigram_headreorientations.mp4**

Same as Supplementary movie 1, but for a trigram (sequence of syllable 2, 23, and 2 again).

**Movie S4. syllable16-1-6_trigram_scrunch-walk-walkandscan.mp4**

Same as Supplementary movie 1, but for a trigram (sequence of syllable 16, 1, and 6).

